# VLab4Mic: prediction of structural resolvability in super-resolution microscopy

**DOI:** 10.64898/2026.06.02.729521

**Authors:** Damián Martínez, Bruno M. Saraiva, Tayla Shakespeare, Mark Bates, Dylan M. Owen, Christophe Leterrier, Mario Del Rosario, Ricardo Henriques

## Abstract

Determining whether a microscopy experiment can resolve a specific feature of a protein assembly remains difficult because researchers must balance imaging modality, labelling strategy, and probe choice. We present VLab4Mic, a simulation platform that predicts structural resolvability before experiments. Starting from atomic models from the PDB or AlphaFold predictions, VLab4Mic places antibodies, nanobodies, chemical linkers, or fluorescent proteins on epitopes, applies stochastic labelling and steric constraints, and generates virtual samples for widefield, confocal, AiryScan, Stimulated Emission Depletion (STED), and Single-Molecule Localisation Microscopy (SMLM). Simulations of nuclear pore complexes agree with experimental data across all five modalities. In case studies, HIV capsid appearance depended strongly on orientation, and STED and SMLM separated domed from flat clathrin lattices where confocal and AiryScan did not. VLab4Mic therefore lets researchers judge which biological questions a given imaging configuration can answer before spending time tuning parameters at the microscope.

## Main

Super-resolution microscopy (SRM) has bridged the gap between cellular imaging and structural biology, enabling visualisation of macromolecular assemblies inside cells (2, 3). Although modern optics can localise a fluorophore with nanometre precision, high optical resolution alone does not guarantee that the underlying biological structure is experimentally distinguishable. Recovering a biologically meaningful feature further depends on probe geometry, linkage error, labelling stochasticity, molecular organisation, and imaging modality. The central challenge is therefore not simply achieving high resolution, but determining whether a given experimental configuration can faithfully preserve biologically relevant structural information, a property we call *structural resolvability*. This captures the ability to detect specific biological features despite stochastic fluorophore binding and the physical dimensions of the probes used, presupposing that the structure is detectable above noise; this sensitivity is governed by photon budget, labelling density, and detector noise (modelled in Supp. Note 2).

Choosing between antibodies, nanobodies, or chemical tags, and matching them to an imaging modality, is usually a matter of costly trial and error. Current simulation tools for SMLM experiments have addressed critical aspects that affect the quality of acquired data such as label geometry, labelling efficiency and fluorophore photophysics (Table S6), and can provide a “ground truth” to benchmark experimental designs and algorithms (4–9). Other simulators instead start from molecular kinetics, for example to test whether transcription-factor clustering reflects genuine condensates or imaging effects (10). While specific studies have used microtubules (11) or nuclear pore complexes (12) with structural detail, general-purpose simulators typically oversimplify the biological sample, representing structures as an idealised geometric shape. This disconnect means that questions critical to structural biologists, such as whether a structural change can be resolved with a primary-secondary antibody pair or how structural integrity affects the ability to measure pore diameter, cannot be reliably answered *in silico*.

Starting from atomic-resolution models, VLab4Mic models how molecular geometry, epitope accessibility, probe architecture, steric constraints, and stochastic labelling together determine whether a biological structure stays experimentally resolvable. This atomic input is essential for super-resolution modalities, and remains informative for diffraction-limited modalities by capturing how crowding and orientation effects influence apparent intensity. By applying physically realistic labelling algorithms across widefield, confocal, AiryScan, STED (13), and SMLM (14–16) modalities, VLab4Mic captures the steric hindrance and binding heterogeneity that dictate structural resolvability.

This framework separates resolvability from sensitivity: resolvability asks whether a specific biological feature, such as a ring radius or a curvature transition, is recoverable from the simulated image, whereas sensitivity governs general detectability above noise. The two are not orthogonal in practice (image-similarity metrics like SSIM remain coupled to noise), but VLab4Mic exposes them as separately controllable inputs (probe geometry and labelling efficiency on the resolvability side; photon budget and detector model on the sensitivity side), enabling biological-feature hypothesis testing rather than mere instrument tuning. Researchers can pose concrete structural questions *in silico* to estimate whether functionally distinct structural states are likely to remain experimentally distinguishable under a proposed experimental setup; for example, users can generate testable predictions regarding whether domed and flat clathrin lattices are structurally resolvable. By predicting how biology, chemistry, and physics shape the final image, VLab4Mic supports experimental design in structural cell biology.

Imaging experiments in VLab4Mic mirror the stages of experimental design: selecting a structure of interest, preparing the sample by selecting a probe, choosing imaging techniques, and analysing results. The workflow (Fig. 1) begins with the selection of an atomic structure from the Protein Data Bank (17) and the specification of a labelling strategy (antibody, nanobody, or chemical linker), which together define target binding sites and probe spatial properties (Fig. 1a). This approach generates a structural model with precise fluorophore positioning, accounting for constraints imposed by both the structure and labelling strategy. The labelling model incorporates stochastic elements such as incomplete binding and structural heterogeneity (Fig. 1b). Multiple independent copies of labelled structures are distributed in three-dimensional space to form a virtual sample. Imaging is then simulated through a forward model that convolves the fluorophore distribution with an *effective post-reconstruction PSF* per modality (wide-field, confocal, AiryScan, STED; modality-specific σ in Table S3), whereas SMLM is instead generated by sampling each emitter from a Gaussian at its localisation precision (see Methods). VLab4Mic therefore produces the image a researcher would obtain *after* any modality-specific reconstruction step, not raw camera frames fed through a reconstruction pipeline (Fig. 1c). A validation framework then systematically compares the simulated images to reference features to quantify structural resolvability (Fig. 1d). The complete computational pipeline is detailed in Fig. S1 (Sup. Video 1).

**Fig. 1.**
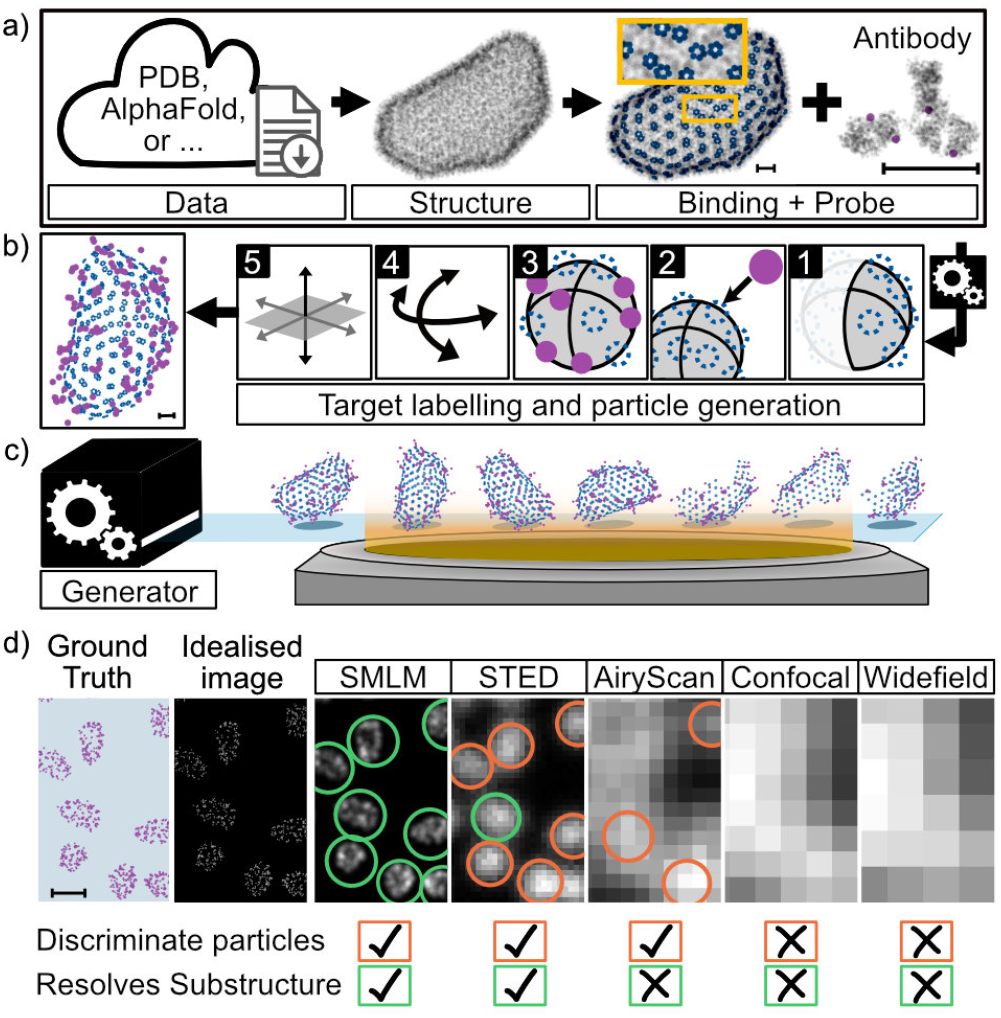
VLab4Mic workflow. **a) Structure and probe selection**. Users can select or provide PDB/CIF structures from databases like PDB or AlphaFold. These files contain atomic coordinates shown in grey. VLab4Mic parses this data to map specific epitope sites in blue. A chosen structural model is used to build a probe. For example, an antibody probe is shown with four conjugated fluorophores in magenta. **b) Epitope labelling**. The platform models the binding of probes to available epitopes. Users can model structure heterogeneity first to restrict available binding sites (step 1). Probes are repositioned toward available epitopes (step 2). Probe constraints may cause incomplete labelling (step 3). The final labelled particle becomes a rigid entity. It can rotate (step 4) and translate (step 5) independently. **c) Virtual sample generation**. Labelling is a stochastic process. A single probe and structure combination can create multiple unique labelled structures. These structures are arranged in space to build a virtual sample. **d) Imaging simulation**. Virtual samples act as the ground truth for imaging simulations. The software simulates modalities like widefield, confocal, AiryScan, STED, and SMLM. Additional pipelines quantify imaging features, such as the ability to resolve substructures or discriminate particles. Scale bars: 10 nm (a, b); 100 nm (d).

Each probe is represented by the spatial distribution of its fluorophores (magenta) relative to the probe structure (grey) and an anchor at the target binding site (Fig. S2), capturing the epitope-to-fluorophore distance that sets the achievable localisation precision (probe specifications in Table S1). For diffraction-limited modalities (widefield, confocal, AiryScan) this geometry effectively reduces to the probe centre, but it becomes important for SMLM and other super-resolution techniques, where probe-induced linkage error sets a floor on the apparent feature size.

Probes are positioned on epitopes by a rigid-body algorithm that orients each along the local epitope normal vector, with a wobbling term modelling linker flexibility. Multiple copies are placed to build a labelled structure (Fig. S8, Fig. S2; algorithm detailed in Methods). Primary-secondary probe pairs are also supported, with secondaries binding the structure-bound primaries under their own independent constraints to capture the larger linkage error of indirect labelling (Fig. 3c, Fig. S6; procedure in SI Methods).

To test the forward model against experimental data, we used NPCs, a widely used nanoscale reference standard for quantitative super-resolution microscopy with experimentally validated dimensions. We generated virtual samples matching the structure, epitope site, probe, and imaging parameters reported by Thevathasan et al. (1) (see SI Methods). Because particle positions were primed from the local maxima of the corresponding experimental images (seeded at the same positions as the real individual NPC structures), this is a consistency check on per-particle appearance rather than a prediction of the field layout (Supp. Note 3). Across all five modalities, from diffraction-limited widefield through super-resolved SMLM, the simulated images reproduce the appearance of the experimental data (Fig. 2). Quantitatively, apparent NPC radii measured by image-based circle-fitting on simulated and experimental SMLM data share the same bimodal distribution, with a primary mode at approximately 52 nm and a secondary mode at approximately 65 nm (Fig. S10). The primary mode found in our radii estimations closely matches the reported Nup96 radius of 53.7 ± 2.1 nm (1). This recovery should be interpreted as a consistency check rather than an independent validation of the NPC radius. The informative result is the secondary mode: our simulated NPCs are identical in every particle by construction, yet an image-based circle fit recovers the same second mode from the simulation as from the experimental images, showing that the 65 nm mode does not reflect a distinct NPC subpopulation (circle-fit procedure in Supp. Note 3).

**Fig. 2.**
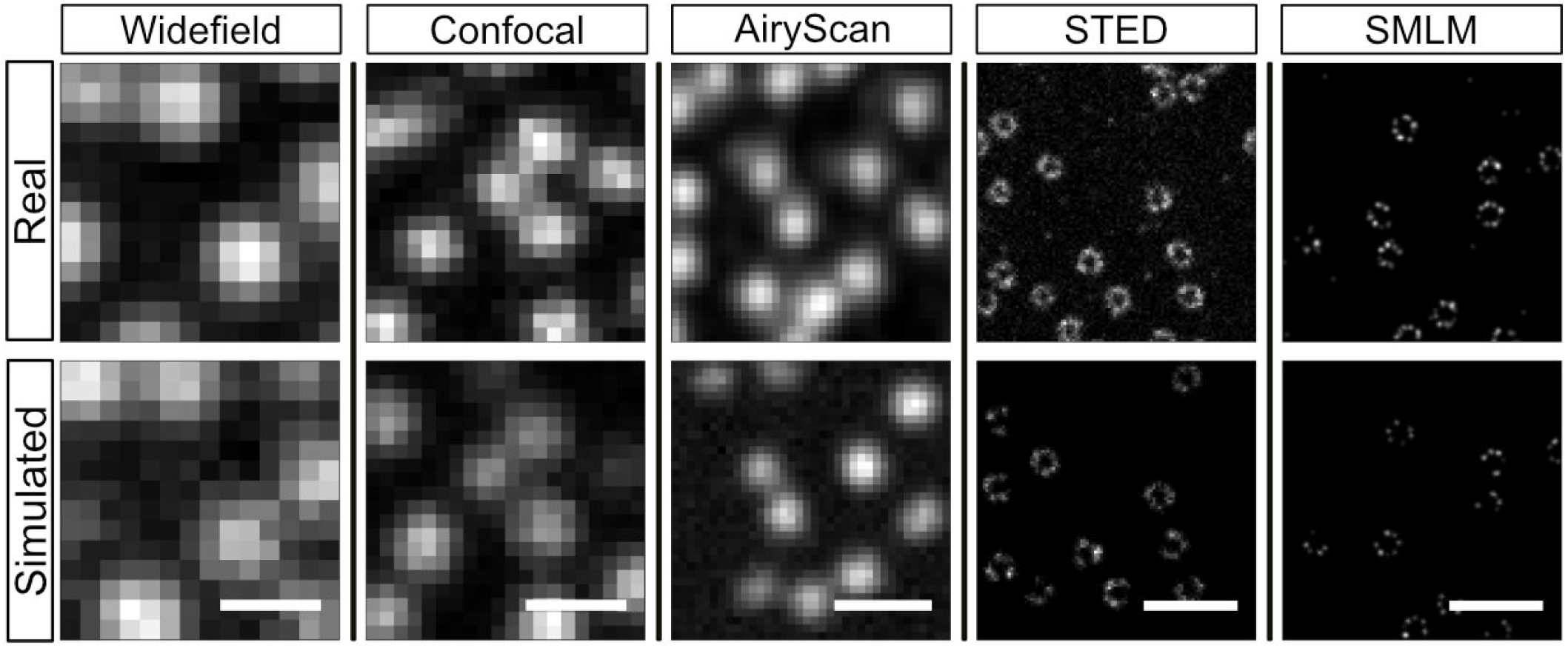
Comparison of VLab4Mic imaging simulations with experimental data. VLab4Mic imaging simulations based on available experimental images of Nuclear Pore Complexes (1). For each modality, the positions of NPCs in the virtual sample were estimated as the local maxima peaks from its corresponding experimental example (See Methods). Scale bars: 500 nm.

VLab4Mic can also sweep parameters automatically: a branching workflow generates all combinations of structural properties (labelling efficiency, structural completeness) and detection characteristics (imaging modality, microscope settings) from user-defined ranges, scoring each against a reference and summarising the space as heat maps (Fig. 3c,d; Sup. Video 2). This replaces manual parameter testing with quantitative metrics for comparing experimental configurations (tested ranges in Table S2; per-cell SSIM/Pearson grid in Fig. 3d).

Virtual samples can be systematically parameterised by particle position relative to the focal plane and by rotational orientation (Fig. 3a,b; Fig. S4; Fig. S5), enabling quantification of how 3D sample geometry affects 2D image formation. This addresses a recurring difficulty in comparing simulated and experimental images: experimental imaging captures random particle orientations and axial positions, whereas conventional simulations often depict ideally oriented structures. By generating distributions of orientations and depths, VLab4Mic produces image ensembles that reflect this experimental heterogeneity.

**Fig. 3.**
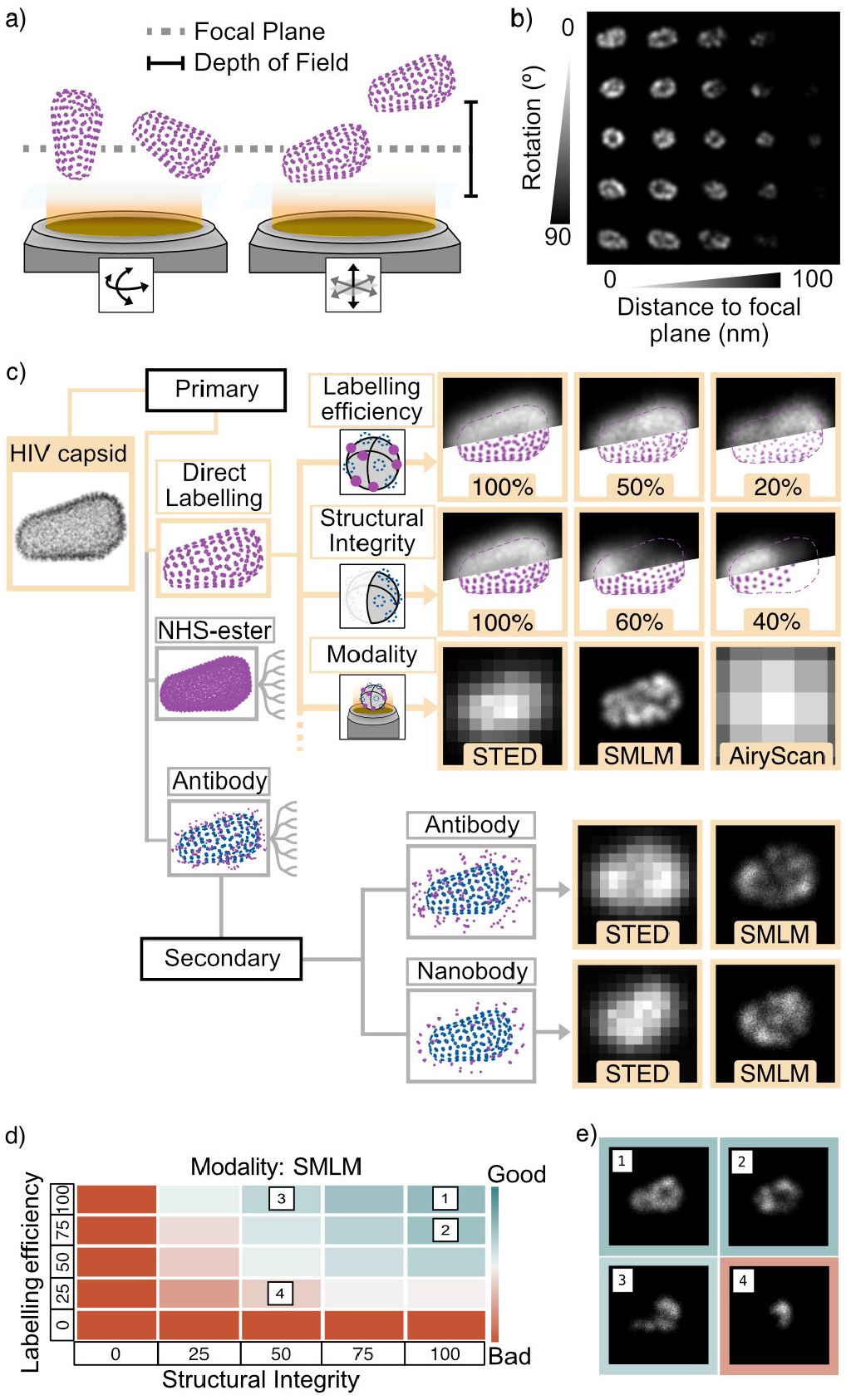
Parameterisation of Virtual Sample, Labelled Structure and Parameter sweeps using HIV capsid model. **a) Virtual sample parameterisation**. Schematic depicting how a labelled particle, such as HIV capsids, can be independently parameterised; for instance, with different orientations or positions relative to focal plane. Magenta dots represent fluorophores. **b) Axial orientation and position sweep**. SMLM image simulation of a virtual sample where a labelled particle’s axial orientation and axial position varies along the X and Y axis, respectively. **c) Parameter branching**. The HIV capsid structure (PDB: 3J3Y, grey dots in atom locations) is labelled with different probes targeting the p24 Capsid protein; epitope sites in blue, fluorophores in magenta. Labelling efficiency (100%, 50%, 20%), structural integrity (100%, 60%, 40%), and imaging modality (STED, SMLM, AiryScan) are varied independently. A dotted line indicates the expected cone shape for structural integrity examples. Secondary labelling is also supported, with primary and secondary probes independently parameterised. **d) Parameter sweep**. VLab4Mic sweeps user-defined parameter ranges (here, labelling efficiency and structural integrity at 5 values from 0–100%) and compares each combination to a reference structure via similarity metrics. High-similarity regions indicate experimentally recoverable states; low-similarity regions indicate loss of structural distinguishability. Results are summarised as a heat map table. **e) Example sweep images**. Representative images from the parameter sweep in d).

Accurate mapping of a structure is limited by the effective labelling efficiency, which is a combination of the number and disposition of the available targets, and the probe binding affinity. The parameter sweep workflow quantifies the tradeoffs between labelling completeness and structural integrity (Fig. 3e), reporting structural resolvability as the Structural Similarity Index (SSIM) and Pearson correlation between each simulated image and a fully-labelled, fully-intact reference (definitions in Supp. Note 3, Eq. 14 and Eq. 15; the tested parameters for HIV-1 capsid sweep are described in Supplementary Notes). We note that this reference is itself simulated and so SSIM/Pearson here measure agreement with the model’s idealised assumptions rather than with biological ground truth; for biological-truth comparisons VLab4Mic supports task-specific metrics (radial distribution analysis for ring structures, Fourier-space periodicity for lattices) discussed below. At high labelling efficiency and complete structural integrity, the simulated cone shape of the HIV-1 capsid recovers the reference (Fig. 3e top left). SSIM declines as either parameter degrades: at 50% structural integrity with complete labelling, the cone outline is preserved while internal density is distorted, and at jointly low labelling efficiency and structural integrity the cone shape is no longer visually discernible (Fig. 3e bottom panels). Both labelling efficiency and structural integrity therefore need weighing when designing an experiment, since either alone can render a key feature unresolvable. Structural integrity modelling algorithms and validation are detailed in Supp. Note 1 and Fig. S3.

VLab4Mic applies to diverse biological structures and labelling strategies (Fig. S13), and makes it straightforward to add custom structural models for resolvability assessment (Fig. 4). As an example, we used ChimeraX (19) to assemble clathrin-coated structures (CCS) as either domed or flat lattices (Fig. 4a), two conformations that coexist on cellular membranes and are implicated in clathrin-mediated endocytosis (18). Because the exact arrangement of triskelia within domes and plaques is not yet known (20), the models shown here were assembled by us as illustrative examples, guided by the PREM reference (Fig. 4b). Each CCS model was equally labelled and simulated under four imaging modalities. VLab4Mic predicts that confocal and AiryScan do not visibly distinguish the two conformations, whereas STED and SMLM resolve them as distinct geometries (Fig. 4).

**Fig. 4.**
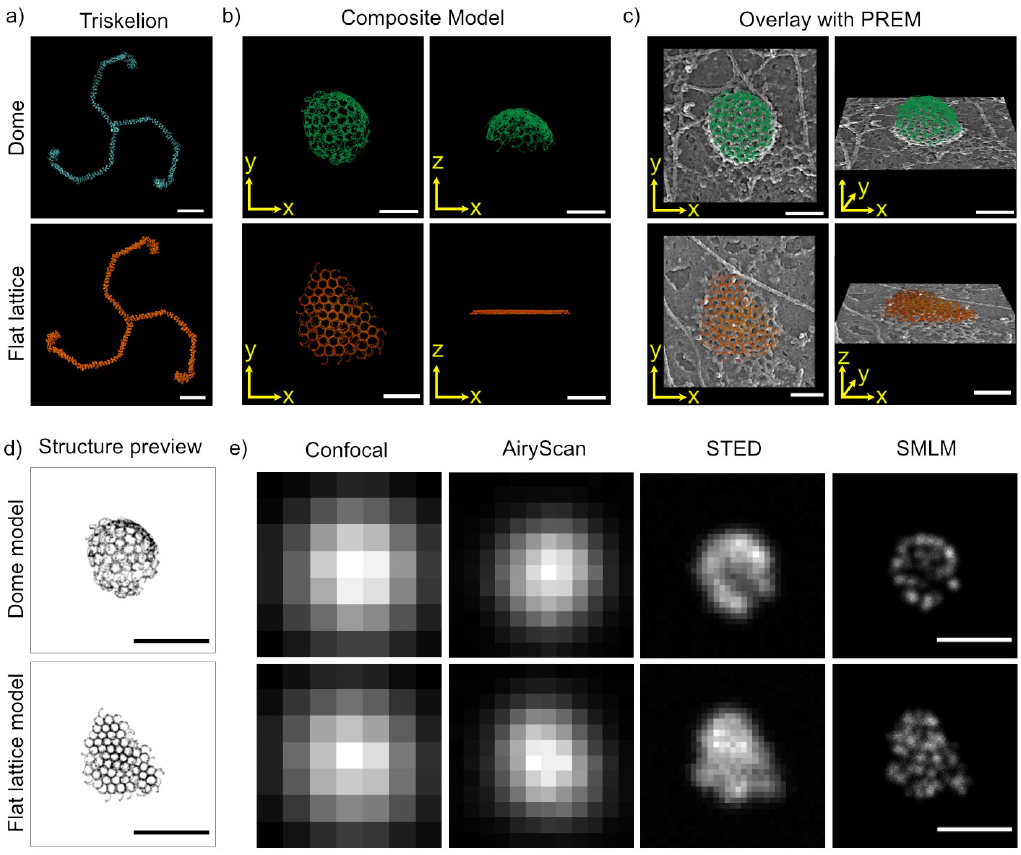
Building and testing custom structural models with ChimeraX and VLab4Mic. Structures assembled in external software can be imported into VLab4Mic to predict how distinct conformational states appear under different imaging modalities. **a) Triskelion models**. Triskelion building blocks used to create composite models were derived from a related clathrin coat assembly (3 copies of the heavy chain from the 3IYV model were selected and arranged to create each triskelion). **b) Composite CCS models**. Clathrin Coated Structures (CCSs) assembled in ChimeraX as a dome (top) or flat lattice (bottom), placing copies of a PDB to match a Platinum Replica Electron Microscopy (PREM) reference (18) (see methods). Models were exported as CIF for VLab4Mic and are available on Zenodo (see Data availability section). **c) Overlay with reference PREM**. Composite models overlaid on the reference PREM image. **d) Structure preview**. Composite models previewed in VLab4Mic. A small fraction of all the atoms is used for visualisation. **e) Simulations across modalities**. VLab4Mic simulations of the dome and flat lattice under four imaging modalities. Both models were labelled with the same antibody probe targeting the heavy chain (see SI Methods). The dome and flat lattice are predicted to be distinguishable in STED and SMLM but not in confocal or AiryScan. Scale bars: 10 nm (a); 100 nm (b, c); 200 nm (d, e).

VLab4Mic also accepts AlphaFold predictions as input (Fig. S11; Sup. Video 3), which is particularly useful for targets lacking experimental structures where resolvability questions arise before crystallography or cryo-EM is attempted. For structures where individual subunits need to be distinguished rather than averaged, VLab4Mic supports chain-specific multichannel labelling (Fig. S12). Using PCNA as a protein nanoruler with the same residue targeted in each chain, VLab4Mic predicts that SMLM with effective *σ* ≈ 2 nm does not resolve the 6 nm inter-site spacing, while a custom SMLM modality with *σ* ≈ 1 nm does; multichannel labelling breaks the trimer’s apparent symmetry and recovers per-chain topology in the simulation even when single-channel resolution is limiting. These examples illustrate how VLab4Mic can predict whether a specific structural question is experimentally tractable with a given imaging modality.

VLab4Mic provides a structure-agnostic forward model that integrates structural biology data with realistic labelling and imaging models, and complements other computational tools (4, 5, 11, 12). Its key advance is the *in silico* capacity to ask whether a structural feature can be resolved before any experiment, testing the resolving power of different probe, labelling and imaging settings.

The model reproduces the second mode of the apparent NPC ring-radius distribution from a population of identical particles, so this mode does not reflect a distinct NPC subpopulation. The framework therefore captures key sources of variability in structural super-resolution imaging (21). Scoring how labelling and acquisition parameters affect structural fidelity also speaks to the wider problem of quantifying imaging artefacts (22, 23). By moving optimisation into a virtual environment, VLab4Mic can reduce trial-and-error and unnecessary acquisition time.

VLab4Mic addresses practical experimental design questions: which modality and labelling strategy can resolve a given feature, how much displacement a primary-secondary antibody pair introduces, or when a design will fail outright, as the orientation-dependent HIV-1 capsid case illustrates (Fig. 3). It is not intended to replace experimental validation, but to guide design toward configurations most likely to preserve biologically meaningful structural information.

To stay tractable across all five modalities, the forward model makes deliberate simplifications: structures are static, point spread functions are analytical, and detector noise follows a single camera-style model (quantum efficiency, stochastic gain, Gaussian readout, and digitisation) applied to every modality (Supp. Note 2). In SMLM, each emitter is rendered at its localisation precision rather than as a series of blinking events; in the diffraction-limited modalities (widefield, confocal, AiryScan), the fluorophores within a single probe fall well inside the blur and are averaged to a single point. The model is therefore most informative when image contrast is governed by particle orientation and molecular crowding rather than by photon statistics.

The scope of these claims is bounded by what the model takes in versus what it produces. VLab4Mic assumes the structure, probe geometry, labelling statistics, and detector parameters are user-specified, and predicts the resulting image and which features survive it; it does not infer unknown structure. A match with an experimental image therefore confirms only that the inputs and the image-formation model are mutually consistent, not that either is biologically correct. The model would be falsified if, for a target with independently known ground truth, the predicted resolvability of a feature disagreed with experiment; closing that loop prospectively would upgrade these consistency checks to predictive validation.

Because VLab4Mic generates synthetic images with known atomic ground truth, it complements curated experimental SMLM resources (24) and to emerging deep-learning analyses that require systematic benchmark generation, including contrastive-embedding methods for nanoscale spatial-omics (25). It integrates with established analysis frameworks (26, 27) and broader deep-learning workflows (28, 29). As super-resolution microscopy moves toward molecular-scale imaging (30–34), frameworks that test whether a structure is not just visible but resolvable will matter more for experimental design and quantitative structural biology.

## ABOUT THIS MANUSCRIPT

This manuscript was prepared using R*χ*iv-Maker v1.22.1 (35).

## DATA AVAILABILITY

Structural models are publicly available from the RCSB Protein Data Bank (https://www.rcsb.org): 7R5K (Nuclear Pore Complex), 3J3Y (HIV-1 capsid), 1XI5 (Clathrin), 3IYV (clathrin heavy chains used for creation of the dome and flat-lattice assemblies in ChimeraX), 8GMO (T4 phage). Experimental NPC images and localisation tables used for the consistency check against published NPC geometry are deposited in BioStudies under accession S-BIAD8 (https://www.ebi.ac.uk/biostudies/bioimages/studies/S-BIAD8) (36), associated with the original publication (1). Supplementary data generated for this manuscript is available through Zenodo (https://doi.org/10.5281/zenodo.20377069). All simulation parameters and configuration files used in this manuscript are provided in the VLab4Mic repository examples directory.

## CODE AVAILABILITY

VLab4Mic source code, installation instructions, API documentation, and example notebooks are available at https://github.com/HenriquesLab/VLab4Mic(v0.1.0) under the MIT licence. VLab4Mic requires Python 3.10-3.13 and runs on macOS, Linux, and Windows. LabConstrictor executables are available at https://github.com/HenriquesLab/LabConstrictor-VLab4Mic. All code used to generate figures and analysis in this manuscript is provided in the repository examples directory.

## AUTHOR CONTRIBUTIONS

**Conceptualisation:** D.M., B.M.S., M.D.R., R.H. **Methodology:** D.M., B.M.S., T.S., M.D.R., R.H. **Software:** D.M., B.M.S., T.S. **Investigation:** D.M., B.M.S., T.S. **Formal analysis:** D.M. **Resources:** B.M.S. **Data curation:** D.M. **Visualisation:** D.M. **Writing - original draft:** D.M., B.M.S., M.D.R., R.H. **Writing - review editing:** D.M., B.M.S., T.S., M.B., D.M.O., C.L., M.D.R., R.H. **Supervision:** B.M.S., M.D.R., R.H. **Project administration:** B.M.S., M.D.R., R.H. **Funding acquisition:** R.H. All authors read and approved the final manuscript.

## ACKNOWLEDGEMENTS

R.H. acknowledges funding from the European Research Council (ERC) under the European Union’s Horizon 2020 research and innovation programme (Self-Driving4DSR, grant agreement No. 101001332). R.H. is further supported by the European Union through Horizon Europe (RT-SuperES, grant agreement No. 101099654); by a European Molecular Biology Organization (EMBO) Installation Grant (EMBO-2020-IG-4734); by a Chan Zuckerberg Initiative Visual Proteomics Imaging award (vpi-0000000044, https://doi.org/10.37921/743590vtudfp); by a joint Wellcome, Chan Zuckerberg Initiative, and Kavli Foundation Essential Open Source Software for Science Cycle 6 award (Wellcome 313383/Z/24/Z; CZI EOSS6-0000000260); and by the La Caixa Foundation (CaixaResearch Health 2025, VirusAwareScopes, HR25-00453). This work was supported by FCT – Fundação para a Ciência e a Tecnologia, I.P., through the MOSTMICRO-ITQB RD Unit (DOI 10.54499/UID/04612/2025, UID/PRR/4612/2025) and the LS4FUTURE Associated Laboratory (DOI 10.54499/LA/P/0087/2020). The Chan Zuckerberg Initiative awards are made through the Chan Zuckerberg Initiative DAF, an advised fund of Silicon Valley Community Foundation. D.M.O. acknowledges funding from the Biotechnology and Biological Sciences Research Council (BBSRC, grant BB/X018644/1). M.D.R acknowledges the Center of Excellence IMMENS funded by the Research Council of Finland (374180). C.L. acknowledges funding from the French National Research Agency (ANR-24-CE13-7996 “MicRON”), and from Excellence Initiative of Aix Marseille University - A ∗ MIDEX (DINoMIR AMIDEX AMX-22-RE-AB-137 to C.L., NeuroMarseille AMX-19-IET-004). We would like to thank the Neuro-Cellular Imaging Service and Nikon Center for Neuro-NanoImaging at INP, with funding from CPER-FEDER (Plateforme NeuroTimone PA0014842), as well as the Institut Neuro-Marseille (AMX-19-IET-004) and the Institut Marseille Imaging (AMX-19-IET-002) for complementary equipment funding from Excellence Initiative of Aix-Marseille University – A ∗ MIDEX, a French “Investissements d’Avenir” program (ANR-11-IDEX-0001). Funded by the European Union. Views and opinions expressed are, however, those of the authors only and do not necessarily reflect those of the European Union or the granting authority. Neither the European Union nor the granting authority can be held responsible for them. The authors thank all members of the Henriques laboratory for helpful discussions, Iván Hidalgo-Cenalmor and Guillaume Jacquemet for their help with the LabConstrictor implementation, and Mike Heilemann for helpful feedback on the manuscript, and Stéphane Vassilopoulos for feedback on the clathrin models.

## EXTENDED AUTHOR INFORMATION

## Methods

### Software implementation and availability

VLab4Mic is implemented in Python 3.10-3.13 and available as an open-source package on GitHub (github.com/HenriquesLab/VLab4Mic) under the MIT licence. The core dependencies include BioPython for PDB/CIF parsing (37), NumPy and SciPy for numerical operations, and standard scientific Python libraries for image processing and visualisation. VLab4Mic provides four usage modes: (1) widget-based Jupyter notebooks requiring no coding, (2) Python API for programmatic access, (3) Google Colab notebooks, and (4) LabConstrictor (38) executables. Zero-installation Google Colab notebooks let researchers run VLab4Mic and optimise protocols without high-end local hardware. VLab4Mic Jupyter notebooks are supported through EZInput widgets for parameter selection (39). All simulations presented in this manuscript can be reproduced using the example scripts and notebooks provided in the repository.

### Structural model generation

We obtain atomic-resolution structures from the RCSB Protein Data Bank (17) in PDB or mmCIF format (structural models used in this study are listed in Table S5). VLab4Mic uses BioPython to parse these files and extract atomic coordinates, chain information, and residue identities (Fig. S7). Users specify target sites using flexible selection syntax that can identify specific residues, chains, or structural motifs.

### Probe positioning

Each probe type (antibody, nanobody, fluorescent protein, or chemical linker) is defined by a structural model showing the spatial distribution of fluorophore positions relative to an anchor point. Probes are placed by a rigid-body algorithm that makes the anchor coincident with the target site and aligns the probe’s principal axis with the local epitope normal (Fig. S8), ensuring physically realistic orientation relative to the protein surface. The normal is calculated via the concentric shell approximation, approximating each site’s local surface normal as an outward-facing vector (Supp. Note 1, Surface Normal Calculation, Eq. 3). A wobbling parameter introduces angular variation by perturbing the normal within a cone of specified opening angle, modelling orientational flexibility and the resulting fluorophore localisation uncertainty (Fig. S2c; Supp. Note 1, Eq. 4). Multiple independent copies of the probe are positioned on the epitopes to generate a labelled structure (Fig. S2d).

### Labelling efficiency and structural integrity

Labelling efficiency is modelled as a Bernoulli process where each epitope site has a probability of being labelled based on the specified efficiency parameter (Supp. Note 1, Eq. 1). Structural integrity is modelled by stochastic removal of clustered subunits prior to positioning probes (Fig. S3; Supp. Note 1, Eq. 5). This structural integrity model should be interpreted as a phenomenological approximation of spatially correlated structural incompleteness rather than a mechanistic representation of biological disassembly. The approach is intended to emulate experimentally relevant perturbations such as partial fixation artefacts, incomplete assembly, lattice collapse, or local molecular loss. The first step uses DBSCAN (40) to cluster epitopes into stable subunits defined by a short-distance parameter. The second step selects a random breaking point, from where neighbouring subunits will be removed. A large-distance clustering parameter limits the distance from which two subunits are considered neighbours. An iterative process removes neighbouring subunits until the number of remaining subunits closely matches the specified structural integrity value (structural integrity clustering algorithms are described in Supp. Note 1).

### Virtual microscope implementation

The imaging simulation treats each fluorophore as a point source emitting photons (41). Image formation is simulated by convolving the effective PSF model of each modality (42) with a discrete version of the fluorophore distribution (Supp. Note 2, Equations Eq. 7 to Eq. 9) and applying a noise model accounting for shot noise, readout noise, and EMCCD gain (Supp. Note 2, Equations Eq. 8 to Eq. 13). For SMLM, each emitter position is also sampled from a Gaussian centred on its true coordinate, with σ set to the modality’s localisation precision, mimicking the accumulation of localisations across frames (Supp. Note 2). The current noise model is calibrated for camera-based wide-field and SMLM detection; consequently, confocal, STED, and AiryScan images should be read as PSF-shape demonstrations rather than as photometrically faithful representations of point-detector chains (such as PMT, APD, or GaAsP). Virtual microscope parameters for all modalities are listed in Table S3.

### Parameter sweep framework

The parameter sweep functionality generates a full factorial design across user-specified parameter ranges. Each parameter combination defines a unique simulation that is executed independently. We score each simulation against a reference, either a ground-truth model or a user-specified target, using structural similarity (SSIM) and Pearson correlation (defined in Supp. Note 3, Equations Eq. 14 and Eq. 15). User-defined metrics can be plugged in as required. Results are aggregated into multidimensional tables that can be visualised as heat maps or line plots.

### Comparison against experimental NPC data

For the NPC consistency check, we used structure 7R5K from the PDB and selected C-terminal residues of Nup96 subunits as target sites. Extended modality-specific parameters for the NPC virtual sample, PCNA site-specific labelling, and the ChimeraX clathrin-coated-structure modelling protocol are detailed in Supp. Note 4. Virtual microscope parameters, detection parameters, including pixel size, field of view, and noise characteristics, were specified to closely match reported experimental values where available (1). Virtual Samples for each modality example were generated by placing multiple copies of the labelled structure at positions corresponding to local maxima in the experimental images, ensuring that the simulated sample closely mimics the spatial distribution of NPCs observed experimentally (Supp. Note 1). For radius measurements, we implemented an image-based circle fit with equal parameters to both simulated and experimental images as described in Supp. Note 4.

## Supplementary Information

**Sup. Table S1.** Probe library and specifications. Specifications of the varying probe models available in VLab4Mic, detailing their local source models, linkage errors (distance), and rotational flexibility (wobble) characteristics. Secondary antibody specifications reflect combined primary+secondary linkage distance. Wobble angles are per-probe values applied at each binding step; the effective wobble of a primary/secondary chain compounds these and can approach 45°(see Structural Modelling and Labelling note below). The geometry and positioning of these probes are detailed in Fig. S2.

| Probe Type | Model ID | Format | Probe Size (30) | Wobble (11) |
| --- | --- | --- | --- | --- |
| Primary Antibody | 1HZH | CIF | ~15-20 nm | 10° |
| Secondary Antibody | 1HZH | CIF | ~20 nm (primary+secondary) | 15° |
| Nanobody | 7MFV | CIF | ~2-4 nm | - |
| GFP-Nanobody | 6XZF | CIF | ~4-6 nm | - |
| NHS-ester | Linker | NA | NA | - |

**Sup. Table S2.** Parameter ranges for sensitivity analysis. Range of values tested during parameter sweep experiments to assess the impact of labelling efficiency, structural integrity, and optical constraints on imaging performance. Type categories: **Probe** (labelling-related parameters), **Structure** (sample integrity), **Acq** (acquisition/imaging settings), **Virtual** (sample construction). Corresponding visualisations for these effects are referenced in the ‘Figure’ column.

| Parameter | Type | Tested Values | Figure |
| --- | --- | --- | --- |
| Labelling efficiency | Probe | 0.2-1.0 | Fig. S3 |
| structural integrity | Structure | 0.3 - 1.0 | Fig. S3 |
| Focal depth | Acq | -200..+200 nm | Fig. S4 |
| Rotation angle | Virtual | 0-180° | Fig. S5 |

**Sup. Table S3.** Optical parameters for virtual microscopes. Default acquisition settings for the different simulated imaging modalities, including Point Spread Function (PSF) dimensions, voxel sizes, and Field of View (FOV). These optical properties govern the image formation process shown in Fig. S1.

| Modality | PSF<sub>xy</sub> $\sigma$ (nm) | PSF<sub>z</sub> $\sigma$ (nm) | Voxel (nm<sup>3</sup>) | Pixel size (nm) |
| --- | --- | --- | --- | --- |
| Widefield | 130 | 400 | 10<sup>3</sup> | 100 |
| Confocal | 100 | 300 | 10<sup>3</sup> | 70 |
| AiryScan | 60 | 230 | 10<sup>3</sup> | 40 |
| STED | 20 | 20 | 5<sup>3</sup> | 15 |
| SMLM | 8 | 8 | 5<sup>3</sup> | 5 |

**Sup. Table S4.**
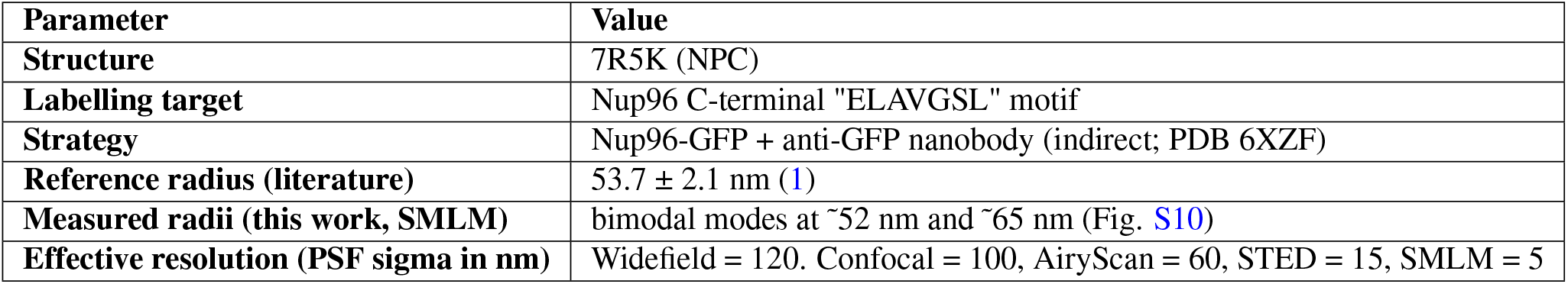
Configuration for validation against experimental data. Parameters used in both the reference experimental dataset (Nup96-GFP labelled with anti-GFP nanobody) and matching VLab4Mic simulations for validation. For the NPC SMLM validation specifically, the STED PSF *σ* deviates from the default in Table S3 (20 nm) and was set to 15 nm to match the experimental dataset; this single-figure override is detailed in the NPC virtual-sample parameterisation methods.

| Parameter | Value |
| --- | --- |
| Structure | 7R5K (NPC) |
| Labelling target | Nup96 C-terminal "ELAVGSL" motif |
| Strategy | Nup96-GFP + anti-GFP nanobody (indirect; PDB 6XZF) |
| Reference radius (literature) | 53.7 ± 2.1 nm (1) |
| Measured radii (this work, SMLM) | bimodal modes at ~52 nm and ~65 nm (Fig. S10) |
| Effective resolution (PSF sigma in nm) | Widefield = 120. Confocal = 100, AiryScan = 60, STED = 15, SMLM = 5 |

**Sup. Table S5.** Macromolecular structural models used for simulations. This table lists the target biological structures, their source identifiers (PDB), physical dimensions, and symmetry properties used as ground truth geometry in VLab4Mic. Clathrin appears at two granularities: PDB 1XI5 is the full auxilin-bound cage used for the diverse-structures and structural-integrity demonstrations (Fig. S13b, Fig. S3c), while PDB 3IYV heavy chains were used to model the triskelion building blocks to assemble the dome and flat-lattice models in ChimeraX for Fig. 4 (see “Clathrin Coated Structures model in ChimeraX” below). The extraction and processing of these structures are illustrated in Fig. S7.

| Structure Name | PDB/CIF ID | Description | Dimensions | Symmetry | Database | Reference |
| --- | --- | --- | --- | --- | --- | --- |
| Nuclear Pore Complex | 7R5K | Human NPC | ~110 nm | 8-fold | RCSB PDB | Mosalaganti et al. 2022 (43) |
| Clathrin Coat (full) | 1XI5 | Auxilin-bound clathrin cage | ~40 nm | Icosahedral | RCSB PDB | Fotin et al. 2004 (44) |
| Clathrin Triskelion | 3IYV | Clathrin D6 coat used to model the dome and flat-lattice CCS (Fig. 4) | ~30 nm | C3 | RCSB PDB | — |
| T4 Bacteriophage | 8GMO | Capsid | ~100 nm | Icosahedral | RCSB PDB | Zhu et al. 2023 (45) |
| HIV-1 Capsid | 3J3Y | Capsid | ~60 nm | Fullerene (conical) | RCSB PDB | Zhao et al. 2013 (46) |

**Sup. Table S6.** VLab4mic features compared against existing microscopy simulation tools.

| Feature | VLab4Mic | SuReSim | SMIS | CIR4MICS | TestSTORM |
| --- | --- | --- | --- | --- | --- |
| **Structure Definition** |  |  |  |  |  |
| Input | PDB ID or PDB/CIF files | Example Pat-terns or surface | Example Patterns | NA | Example pat-terns or surface |
| Macromolecular complex incompleteness | Yes | No | No | No | No |
| **Epitopes** |  |  |  |  |  |
| Epitope positioning | Parsing atomic coordinates from PDB/CIF | Sampling on model | Sampling on model or image | Pre-set positions for NPCs | Sampling on model |
| Multiple epitope identities on structure | Yes | No | No | Up to six labels | No |
| User-defined epitope sites (sequence, residue, etc.) | Yes | No | No | No | No |
| **Labelling and probes** |  |  |  |  |  |
| Direct labelling | Yes | Yes | Yes | Yes | Yes |
| Site-specific labelling | Yes | No | No | No | No |
| Labelling-efficiency | Yes | Yes | Yes | Yes | No |
| Probe size | Yes | Yes | Yes | Yes | Yes |
| Probe orientation | Yes | Yes | Yes | No | Yes |
| Probe structure from PDB (Antibody, GFP, SNAP-tag, etc..) | Yes | No | No | No | No |
| **Fluorophores and photophysics** |  |  |  |  |  |
| Multiple fluorophores | Yes | Yes | Yes | NA | Yes |
| Advanced photophysics (photoconversion, FP-maturation, spectral responses) | No | No | Yes | NA | Polarisation, Cross-talk |
| **Sample and imaging modalities** |  |  |  |  |  |
| Virtual sample from experimental data | Yes | No | No | No | No |
| Expansion Microscopy | Yes | No | No | Yes | No |
| Imaging modalities | SMLM, STED, AiryScan, Confo-cal, Widefield | SMLM | Widefield | No | SMLM |
| Custom modalities | Yes | No | No | No | No |
| 3D-PSF | Yes | Yes | Yes | No | Yes |
| Multicolour imaging | Yes | No | Yes | No | Yes |
| **Miscellaneous** |  |  |  |  |  |
| Parameter sweep | Yes | No | No | No |  |
| Software availability | Python library. Jupyter Note-books. | Java com-piled (7) | Matlab (4) | Python (12) | Matlab (6) |

## Supp. Note 1: Structural Modelling and Labelling

The foundation of VLab4Mic’s simulation capability lies in its ability to generate realistic, physically plausible models of labelled structures. Unlike simple point-source models often used in microscopy simulations, VLab4Mic begins with the ground-truth atomic coordinates parsed from PDB or CIF files. This allows for the definition of target sites (epitopes) with atomic precision, based on user specifications such as specific amino acid residues or peptide sequences

### Structure parsing and target sites

VLab4Mic leverages PDB/CIF models from databases, such as the RCSB Protein Data Bank (RCSB PDB) or AlphaFold, for accurate modelling of probe placing. By starting from the actual molecular geometry (Fig. S7a), the simulation can accurately capture the spatial relationships and constraints that exist in real biological samples. VLab4Mic creates a structural model comprising the atoms within the model and methods to retrieve target sites and associated geometrical properties. We use BioPython parsing tools to read the atoms in the file. However, some assemblies for macromolecular complexes are defined as a single asymmetric unit (highlighted in blue) without explicit definition of the whole complex. For these cases, VLab4Mic constructs the complete assembly model (grey) from the assembly operations in the structure file, as is the case for the Nuclear Pore Complex (NPC) (Fig. S7a Top). From these structures, specific locations on the macromolecular structure are defined as Target sites. Parsing of these sites is specific to the structure and independent of the probe model, allowing for flexible combinations of targets and probes. We consider 2 main types of target definition, which are compatible with PDB/CIF models (Fig. S7b). i. Protein sequence: an amino acid sequence that is compared against all peptide sequences in the PDB/mmCIF. Only exact matches are retrieved as epitopes. The position of the epitope is calculated as the average coordinate from the atoms that belong to that peptide sequence. ii. Residue: specified by a 3-letter code. VLab4Mic iterates over all protein chains and retrieves all instances of the Residue, based on its alpha carbon. The position of the epitope is defined as the coordinate of the alpha carbon atom. VLab4Mic allows for multiple probes to be defined for a single structure, each with its own target specification (Fig. S7c). Additionally, the residue-based target allows for site-specific labelling, where the user can define exact residue numbers to be targeted and the protein chain within the model Fig. S12.

### Probe modelling

A probe describes the position of the fluorophores in 3D space relative to a structural model, such as an antibody or GFP.

- **Fluorophore positions**. The exact positions of the fluorophores can be parsed from the atomic structure of the probe model or as the chromophore coordinates of fluorescent proteins. Common conjugation sites for antibodies, such as lysine residues, can be used to define these positions.
- **Degree of labelling (DoL)**. To capture the variability in the number of fluorophores attached to a probe, VLab4Mic allows users to specify a degree of labelling (DoL) parameter. This parameter defines the average number of fluorophores per probe molecule, and the actual number of fluorophores for each probe instance is sampled from a Poisson distribution with mean equal to the DoL. This allows for realistic variability in fluorophore density across different probes, which can impact the brightness and localisation precision in the simulated images.
- **Probe geometry**. The spatial arrangement of fluorophores on the probe is determined by the probe model. Besides the fluorophore positions, the probe model also defines the paratope site (the site with affinity for the epitope in the structure) and an internal axis that defines the orientation of the probe relative to the target.
- **Distance to epitope**. The distance from the probe paratope to the epitope site can be adjusted with an offset parameter that extends the probe anchor site along the probe axis prior to any probe positioning.

### Probe Positioning and Orientation

- **Stochastic model for structure labelling**. In experimental fluorescence microscopy, it is theoretically impossible for every available epitope to be labelled. Factors such as antibody affinity, incomplete staining protocols, and competition for binding sites result in fractional labelling efficiencies. VLab4Mic simulates this realistic variability through stochastic labelling where each potential epitope site is independently labelled with a probe. Labelling is modelled as an iterative process where each epitope site is evaluated for successful labelling (Bernoulli trial) and then subjected to steric constraints based on the probes already placed.
- **Bernoulli Labelling**. For a given target structure, we assume that each potential epitope site *i* is independent. This assumption simplifies accessibility effects, cooperative binding phenomena, and local environmental heterogeneity, which are not explicitly modelled in the current implementation. The probability of any specific site being successfully labelled is defined by the efficiency parameter *ε*_*label*_. This is mathematically implemented as a Bernoulli trial for each site (Eq. 1):

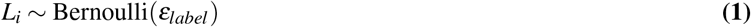

where *L*_*i*_ = 1 indicates a successful label and *L*_*i*_ = 0 indicates an unlabelled site. Per-site independence is assumed; the steric clash term below enforces a minimum spacing between probes but does not encode cooperativity or accessibility-dependent affinity. The effect of varying labelling efficiency on the resulting fluorophore distribution is shown in Fig. S9a.

Once a site is determined to be labelled, the position of the fluorophore relative to the target epitope is not random. Probes are modelled as rigid bodies anchored to the target site, and their spatial orientation is determined by the local surface geometry of the macromolecule. This correctly models the “linkage error” (the distance between the target protein and the actual emitter) which is a critical factor in super-resolution accuracy (47).

- **Steric Constraint Sampling**. Physical probes, such as antibodies or even small organic dyes, occupy physical volume. In dense epitope regions, it is physically impossible for two probes to occupy the same space or for a probe to bind if its footprint is obstructed by neighbours. To account for these steric clashes, VLab4Mic enforces a minimum distance constraint when placing probes in epitope sites. This minimum distance is estimated as the maximum pairwise-length from the atoms within the probe model. A label at site *i* is only permitted if it satisfies a minimum distance *d*_*threshold*_ from all other successfully labelled sites (Fig. S9b). The final distribution of fluorophores reflects the physical exclusion volume of the probes and their efficiency of labelling an epitope.
- **Probe Orientation**. The probe’s principal axis is aligned with the vector normal to the protein surface at the binding site. The final coordinates of the fluorophores is calculated by treating the probe as a rigid body and performing a rotation and translation operation to align the probe axis with the epitope normal vector. We use the Rodrigues’ rotation formula (Eq. 2) to compute this transformation:

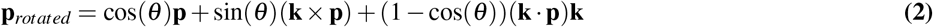

where **p** is the fluorophore coordinate, *θ* is the angle between the probe’s internal axis and the epitope normal, and **k** is the unit vector along the rotation axis defined by the cross product of these two vectors.
- **Surface Normal Calculation**. To orient the probe correctly, the simulation must determine “which way is up” relative to the protein surface. We approximate the local surface normal **N**_*i*_ at target site *i* by comparing the target position on the original structure to a concentric, scaled-down version of the structure (scaling factor *s <* 1, default *s* = 0.95, centroid preserved). The vector from the scaled-down position back to the original provides an outward-facing normal estimate (Fig. S8, Eq. 3):

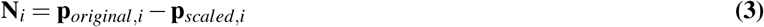

This concentric-scaling approximation is reliable for convex target distributions (e.g., outer rings of NPC, capsid exterior), but degrades on locally concave or planar geometries (e.g., flat clathrin lattices, inward-facing epitopes); for those cases users can supply pre-computed normals or substitute a local-neighbourhood method (e.g., PCA over neighbouring atoms).

- **Angular Wobble**. While probes are modelled as rigid bodies, the linkers connecting them (e.g., antibody hinges or flexible chemical linkers) introduce rotational freedom. This flexibility, or “wobble,” effectively blurs the fluorophore’s position over time, degrading localisation precision (48). The wobble parameter should therefore be interpreted as an effective orientational uncertainty rather than a direct physical measurement of probe dynamics. VLab4Mic models this by introducing random angular perturbations to the probe’s ideal orientation, sampled to be isotropic over the spherical cap defined by a user-specified maximum half-angle *θ*_*max*_ (Eq. 4):

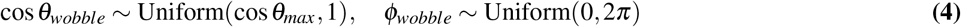

The cos *θ* parameterisation ensures uniform sampling over solid angle within the cone, rather than concentrating samples near the pole.

To accurately model linkage error, we recommend users approximate wobble cones based on the probe’s mechanical properties. For instance, use tighter constraints for nanobodies ( 10°) compared to the flexible hinge regions of primary/secondary antibody complexes ( 45°). This allows the user to simulate the specific localisation uncertainty introduced by their chosen labelling strategy.

### Primary and Secondary probe labelling

To model the commonly used primary and secondary antibody labelling strategy, VLab4Mic allows users to define a two-step labelling process. Primary probes are placed on the target structure using the stochastic labelling and probe positioning methods described above (Fig. S6a). In the second step, secondary antibodies are then placed on the already positioned primary probes (Fig. S6b). The target site for the secondary antibody is defined as an epitope on the primary antibody model. The placement of secondary antibodies is also subject to steric constraints, ensuring that they do not overlap with each other or with the primary antibodies. When secondary labelling is used, the final fluorophore positions are only the fluorophores on the secondary antibodies.

### Structural integrity

Real biological samples are rarely perfect. They exhibit structural heterogeneity, damage from fixation, incomplete assembly or epitope regions inaccessible to the probes. To test the robustness of imaging methods against such imperfections, VLab4Mic explicitly models variations of structural integrity.

- **Cluster-Based structural integrity (DBSCAN)**. Often, damage occurs in larger, contiguous patches, for example, a missing sector of a nuclear pore complex ring. To emulate this large-scale structural damage, we employ Density-Based Spatial Clustering of Applications with Noise(40) (DBSCAN). The first step groups epitopes into stable subunits defined by a short-distance cluster parameter (Eq. 5). We note that this is a phenomenological abstraction rather than a biologically validated damage model; the resulting degradation morphology is one of several plausible regimes, and users may substitute alternative damage rules (for example, per-subunit dropout, patterned ablation, or accessibility-weighted removal) by replacing the cluster-selection routine.

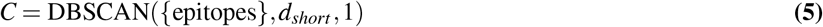

where *d*_*short*_ defines the neighbourhood radius for epitopes to cluster into a subunit. Each subunit is then represented by the average coordinate of the epitopes it contains. Then, a K-d Tree is built for these subunits, so that a distance parameter *d*_*large*_ is used to query neighbouring subunits.

Once grouped, a subunit is selected at random as the breaking point from which an iterative sample process will remove neighbouring subunits. On each iteration, the breaking point is removed and a random subset of its neighbouring subunits will also be removed. On the next iteration a new breaking point is selected from the subunits that are already removed. A large-distance parameter limits the distance from which two subunits are considered neighbours. This process of removal of subunits is repeated until the number of epitopes in the remaining subunits closely matches the specified structural integrity value.

Simulating clustered integrity loss enables researchers to train and validate analysis pipelines, ensuring they can robustly discriminate between structural integrity artefacts and genuine structural gaps in macromolecular assemblies.

### Virtual Sample Construction

A virtual sample consists of multiple independent copies of these labelled structures distributed in 3D space.

- **Labelled Structure Placement**. Labelled structures can be placed within the defined sample volume *V*_*sample*_ by specifying their positions in 3D or by random placing. Crucially, this placement is subject to a non-overlap constraint to prevent physical intersections between distinct molecular complexes (Eq. 6):

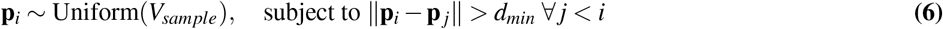
- **Labelled Structure Orientation**. The orientation of each structure is defined by an orientation vector; when reorienting a labelled structure, its central axis, and the rest of the structure, is aligned with the orientation vector. The labelled structure orientation can be defined by the user or randomly assigned. If a random orientation is assigned, the orientation vector is sampled uniformly from the unit sphere, ensuring an isotropic distribution of orientations across the sample. However, users can also specify a bias in the orientation distribution by specifying a set of rotation angles (Euler angles).
- **Labelled Structure planar rotation**. Finally, a planar rotation can be applied to the labelled structure, which is a rotation around the central axis of the structure. This is particularly useful for structures with a defined central axis, such as the NPC, where the planar rotation will rotate the structure around its central axis.

### Virtual Sample priming from input image

VLab4Mic allows users to “prime” a virtual sample with positions derived from a 2D image. VLab4Mic supports two methods for extracting particle coordinates from an input image: (1) local maxima detection and (2) random sampling from binary masks. The resulting virtual sample dimensions are calculated to match the physical dimensions of the input image given its pixel size.

- **Local maxima detection**. This method identifies local intensity peaks in the input image, which are assumed to correspond to individual particles or structures. The coordinates of these peaks are extracted and used as the positions for placing labelled structures in the virtual sample.
- **Random sampling from binary masks**. A binary mask delineates regions of interest in the image where particles are likely to be located. The selection of particle positions is done iteratively by randomly sampling coordinates within the masked regions, constrained by a minimum distance to ensure non-overlapping placement of structures that can be derived by the labelled structure dimensions or a user-defined parameter.

## Supp. Note 2: Image Formation and Photophysics

Once a virtual sample is generated, VLab4Mic image formation process follows a forward model based on an effective PSF model per modality. This effectively acts as a ‘virtual microscope,’ translating the exact 3D coordinates of fluorophores into pixelated images. This process involves modelling the diffraction-limited optics of the system, the photophysical properties of the fluorophores, and the stochastic noise introduced by the detector.

### Point Spread Functions

The primary determinant of image resolution in any optical system is the Point Spread Function (PSF), which describes the system’s response to a point source. Image formation is mathematically modelled as the convolution of the discrete fluorophore distribution with this PSF.

- **3D Elliptical Gaussian PSF**. We approximate the effective PSF of a modality as a 3D elliptical Gaussian distribution where lateral and axial width can be parameterised independently (Eq. 7):

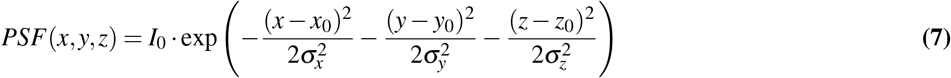

where *σ*_*x*_, *σ*_*y*_, *σ*_*z*_ define the lateral and axial resolution widths, and (*x*_0_, *y*_0_, *z*_0_) is the PSF centre coordinate.

The PSF is a discrete 3D array with sampling rate of 10 nm in each dimension for conventional modalities, while for SMLM and STED, the sampling rate is 5 nm. Users can adjust this sampling rate to balance accuracy and computational efficiency.

### Fluorophore Photophysics

We model the brightness of each fluorophore by defining the average photon emission rate per unit time when the fluorophore is in the ‘On’ state. VLab4Mic models the stochastic behaviour of photon arrival as a Poisson process, which is a common assumption in fluorescence microscopy.

### Scope of the photophysical model

Photon-emission rates and integration times drive shot noise and set the per-emitter localisation count for SMLM (see *Simulation of localisation microscopy modalities* below). VLab4Mic does **not** explicitly model fluorophore photoswitching kinetics (*k*_*on*_, *k*_*o f f*_, dark-state lifetimes, photobleaching) and does not derive blinking duty cycles or exhaustion from first principles. The forward model targets the *effective post-reconstruction image* a researcher would obtain after any modality-specific reconstruction step, rather than simulating raw camera frames followed by a reconstruction pipeline.

- **Photon Emission (Shot Noise)**. For each emitting fluorophore, the number of photons detected during an integration time Δ*t* is a random variable drawn from a Poisson distribution with mean *λ*_*photons*_ · Δ*t*, where *λ*_*photons*_ is the average photon emission rate (Eq. 8). This captures the inherent shot noise in fluorescence imaging, which arises from the discrete nature of photon emission and detection:

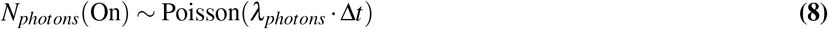

where *λ*_*photons*_ is the average photon emission rate.

### Virtual sample discretization

To perform convolution, the continuous fluorophore distribution must be discretised into a 3D grid of voxels. The sampling rate of the virtual sample is defined by the sampling rate of the PSF of the imaging modality. Before convolution, the number of photons emitted by each fluorophore is calculated based on the photophysical model and integration time (exposure time), so that each voxel contains the total photon count from all fluorophores within it.

### Simulation of localisation microscopy modalities

To simulate the effect of an emitter being localised multiple times across different frames, as is the case in SMLM, we include a previous step where each fluorophore exact position is used to sample new localisation positions from a Gaussian distribution centred on the fluorophore position and with a standard deviation defined by the localisation precision of the modality. An average number of localisations per fluorophore can be defined by the user, and the actual number of localisations for each fluorophore is sampled from a Poisson distribution with mean equal to this parameter. Lateral uncertainty and axial uncertainty are sampled independently. The final image is the result of convolving the PSF of the imaging modality with sampled localisation positions. This localisation simulation is set as default for SMLM, but can be disabled to simulate a SMLM modality without localisation uncertainty. The Poisson per-emitter localisation count is appropriate for PAINT-type acquisitions, in which transient binding events accumulate independently; PALM/STORM-style exponential per-site distributions, where switching of a fixed fluorophore until photobleaching produces an exponential tail of localisations, are not modelled.

### Image formation via convolution and binning

To obtain the final image, the discretised fluorophore distribution is convolved with the PSF of the imaging modality, integrated in the axial dimension and then binned to the pixel size of the detector.

- **Convolution of virtual sample and PSF**. The interaction between the optical system and the sample is linear and shift-invariant. Therefore, the final intensity distribution *I*(*x, y, z*) is computed via Fast Fourier Transform (FFT) convolution of the fluorophore spatial distribution *F* and the system PSF (Eq. 9):

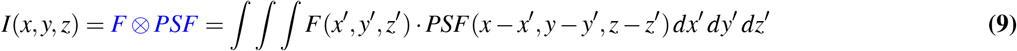

This results in a 3D intensity array with the same sampling rate as the PSF.

- **Depth of Field and Out-of-Focus Fluorophores**. The intensity array is then projected onto a 2D plane by summing along the axial dimension with respect to the focal plane in the sample. The depth of field of the microscope restricts the planes above and below the focal plane that contribute to the image. This projection simulates the depth of field of the microscope, where fluorophores that are out of focus contribute to the image with a broadened PSF in the axial dimension.
- **Pixel Binning**. Finally, the 2D intensity distribution is binned (with addition) to match the pixel size of the detector. This step simulates the finite spatial sampling of the camera, where each pixel integrates the signal from a defined area of the sample. The resulting 2D array represents the ideal, noise-free image formed by the virtual microscope.

### Detector Noise Models

The final stage of image formation occurs at the detector, where photons are converted into digital signals. VLab4Mic simulates detector characteristics by including quantum efficiency, amplification, and electronic readout noise pixel-wise. A single camera-style model (quantum efficiency × stochastic gain × Gaussian readout × digitisation, see equations below) is applied uniformly across modalities. This matches camera-based acquisitions (widefield, SMLM); point-detector modalities (confocal, STED, AiryScan) whose physical detectors (PMT, APD, GaAsP) follow Poisson + dark-count statistics are not modelled at the detector level, and their outputs should be read as PSF-shape and spatial-arrangement demonstrations.

- **Detection Efficiency**. Not every photon hitting the detector generates a signal. The probability of an incident photon being converted into a photoelectron is determined by the detector’s Quantum Efficiency (QE). This loss is modelled as a Binomial process (Eq. 10):

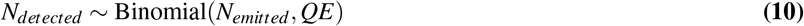

where *N*_*emitted*_ is the number of photons in the noise free image and *QE* is the quantum efficiency of the detector.
- **Electron Multiplication**. Detected photons are amplified before readout. This stochastic amplification process introduces additional noise and is modelled using a Gamma distribution (Eq. 11):

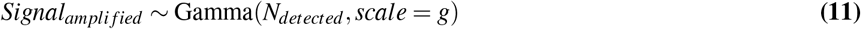

where *g* is the gain factor.
- **Readout Noise and Baselevel**. Finally, Gaussian readout noise, which is independent of the signal level is added to the amplified signal (Eq. 12). A constant baseline offset (BaseLevel) can also be added which is usually done to prevent negative pixel values:

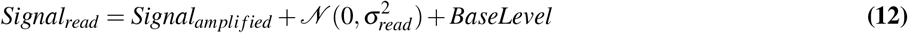
- **Digitisation**. The analogue voltage signal resulting from detection, amplification and readout is then quantised into discrete analogue-to-digital units (ADU), completing the image formation process (Eq. 13):

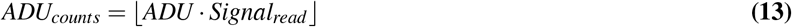

### Multicolour Imaging

VLab4Mic supports multicolour imaging by allowing users to define multiple probes with distinct fluorophores. Each fluorophore will generate a separate image channel for the same modality parameters. The final output is a multi-channel image stack that can be analysed using standard multicolour microscopy techniques.

## Supp. Note 3: Parameter sweep and validation

This section outlines the comprehensive workflow employed by VLab4Mic for systematic parameter exploration and the quantitative metrics used to validate the accuracy of the simulations against experimental ground truth.

### VLab4Mic Computational Pipeline

The overall simulation pipeline integrates the structural modelling and image formation components into a cohesive workflow. As illustrated in Fig. S1, the process begins with atomic structure parsing to define the ground truth geometry. This is followed by the generation of a “virtual sample” (a populated 3D volume of labelled particles). Finally, this virtual sample is effectively “imaged” by the forward model to produce multi-modality synthetic data (Widefield, Confocal, STED, SMLM) from the exact same underlying sample definition.

### Parameter Sweep Workflow

A robust validation framework requires more than single-instance comparisons. To systematically explore the impact of experimental variables on imaging outcomes, VLab4Mic implements an automated parameter sweep generator. This tool creates a factorial design of simulation parameters, varying factors such as labelling efficiency, probe type, noise levels, and optical settings. It then executes batch simulations for each unique combination. This facilitates comprehensive sensitivity analysis, allowing researchers to determine which experimental parameters are most critical for resolving specific structural features.

### Image Similarity Metrics

To objectively quantify the agreement between simulated images (*I*_*sim*_) and experimental references (*I*_*exp*_), we employ standard image quality metrics.

- **Structural Similarity Index (SSIM)**. The SSIM is a perception-based model that considers image degradation as perceived change in structural information. It measures similarity based on three comparisons: luminance, contrast, and structure. Unlike simple mean squared error, SSIM is highly sensitive to the spatial structure of local image patterns, making it ideal for microscopy validation (Eq. 14):

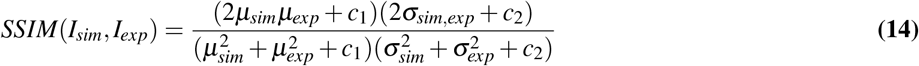

where *µ* and *σ* represent the local mean and variance, and *c*_1_, *c*_2_ are stabilising constants.
- **Pearson Correlation**. The Pearson correlation coefficient (*ρ*) evaluates the linear correlation between pixel intensities of the two images. It provides a measure of how well the intensity variations in the simulation match those in the experiment, independent of absolute brightness scaling (Eq. 15):

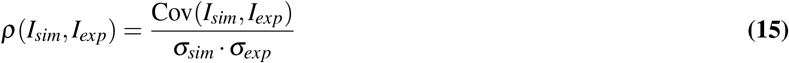

### Comparison Methodology for parameter sweep

Direct comparison of simulated and reference images is often hindered by the random placement of particles in experiments. To address this, VLab4Mic implements a “priming” function for validation. This function extracts the exact particle coordinates from the reference sample. It then uses these coordinates to position the virtual structures in the simulation. This ensures that the spatial arrangement of the simulated sample matches the reference image, allowing for accurate computation of pixel-wise similarity metrics like SSIM.

### Image masks

To further refine the comparison, we use the priming positions to generate a binary mask to limit the analysis to regions where particles are present. By applying this mask to both the reference and simulated images before computing similarity metrics, we focus the validation on relevant regions, reducing the influence of background noise and empty space.

## Supp. Note 4: Extended Methods

### Virtual Sample parameterisation of Nuclear Pore Complexes

Virtual Samples for NPC simulations based on experimental data (1) were generated by positioning independent copies of the NPC labelled with Nup96-GFP in a 1.5 µm x 1.5 µm FOV. We used the PDB 7R5K as model for NPC (Table S4). The probe was based on a GFP with nanobody (PDB: 6XZF), with degree of labelling of 2, epitope defined as the peptide sequence “ELAVGSL” and labelling efficiency of 0.4. The coordinates on where to place the NPCs were obtained by extracting the local maxima peaks with modality-specific image parameters. Widefield: pixelsize = 100 nm, sigma = 0.5, min distance = 100 nm, threshold = 0. Confocal: pixelsize = 70 nm, sigma = 1, min distance = 70 nm, threshold = 400. AiryScan: pixelsize = 40 nm, sigma = 0.5, min distance = 40 nm, threshold = 100. STED: pixelsize = 15 nm, sigma = 3, min distance = 90 nm, threshold = 3. SMLM: pixelsize = 5 nm, sigma = 6, min distance = 120 nm, threshold = 10. Experimental data was obtained from available datasets (36). PSF width for Widefield was set as the mean PSF width measured on the localisation table from the experimental data, and the effective SMLM PSF as the mean localisation precision reported on the same table. PSF widths (sigma) were set as follows: Confocal = 100 nm, AiryScan = 60 nm, STED = 15 nm.

### NPC radii quantification

VLab4Mic includes methods to allow users to prime a virtual sample with positions derived from an experimentally acquired image. We used this functionality to create equivalent simulated images that allow us to reduce the bias of structure placement within a virtual sample for image comparison (Fig. S10). We then measure the apparent radii of NPCs on both simulated and real data to assess if VLab4Mic simulations allow us to recapitulate reported geometrical descriptors of the NPC. Figure (Fig. S10a) shows an example of the STED imaging of NPCs and a VLab4Mic equivalent image simulation for the same modality; the right panel shows their respective histograms for measured radii on each image using the same parameters. Figure (Fig. S10b) shows a similar example but with SMLM images.

Two methods can be used to estimate an NPC radius from SMLM data. Thevathasan et al. (1) fit a circle directly to the table of single-molecule localisation coordinates (a localisation-table fit), and report R = 53.7 ± 2.1 nm. We instead apply circle-fitting to the rendered SMLM images, which is the operation available to a user who has access only to the reconstructed image and not to the underlying localisation table. Under this image-based fitting we observe a bimodal histogram of fitted radii with a primary mode around 52 nm (matching the localisation-table radius reported by Thevathasan) and a secondary mode around 65 nm. Both modes are recovered by VLab4Mic on the simulated equivalent dataset, even though every NPC in the simulation is identical by construction. This indicates the secondary mode is associated with the image-rendering and image-based fitting steps rather than with biological variation in the sample. The agreement between the simulated and experimental distributions supports the use of VLab4Mic to recapitulate image-based quantitative descriptions of the experimental data.

### PCNA site-specific labelling

VLab4Mic allows for site-specific labelling by defining exact residue numbers to be targeted within a protein chain. This feature is particularly useful for simulating experiments where precise labelling of specific amino acids is required, such as labelling of the Proliferating Cell Nuclear Antigen (PCNA), to serve as a protein-based nanoruler (49). Here, the residue SER-186 within each PCNA monomer is targeted, and the expected geometrical distribution of the fluorophores is a triangle of 6nm separation between each target. VLab4Mic accurately models this site-specific labelling (Fig. S12a), where the fluorophore positions (magenta) are shown relative to the PCNA structure (grey). The distances between the fluorophores in our model closely match the expected 6 nm separation. Furthermore, VLab4Mic SMLM simulations with typical resolution parameters (Fig. S12a) do not resolve the three sites as previously reported (49). However, VLab4Mic custom modalities enable the user to explore the conditions required to resolve these closely spaced epitopes: at an effective resolution of 2 nm the three sites remain unresolved as a single feature, whereas at 1 nm they are clearly separated (Fig. S12a).

### Clathrin Coated Structures model in ChimeraX and target selection for VLab4Mic

Dome and flat clathrin coated structures were modelled in ChimeraX by manually placing copies of triskelion derived from 3IYV. To guide the placement of the triskelions, an image of a clathrin coated structure was loaded in the same session. The triskelions were placed and reoriented manually. The resulting model was exported as a PDB file and used as input for VLab4Mic. For the probes to target both models, we chose an arbitrary short sequence (“ATETQ”) that could be parsed from both heavy chains (CHC) as proxy for reported targets for probing CCS ((18)).

### Parameter sweep with HIV-1 capsid

Parameters for HIV-1 capsid sweep were chosen as follows. Structure: “3J3Y”. Probe template: “anti-p24_primary_antibody_HIV”. Number of sweep repetitions: 10. Labelling efficiency and structural integrity parameters: min = 0, max= 1, steps = 0.25. Structural integrity distances were: small cluster = 20; large cluster = 100. Exposure time = 0.001. All other parameters were used as default.

### Interdependence of resolvability conditions

The minimal conditions to achieve structural resolvability are not independent: probe geometry, labelling efficiency, structural integrity, and modality resolution interact, so no single parameter can be optimised in isolation. For instance, a larger probe (a primary/secondary antibody pair rather than a nanobody) increases linkage error, and recovering the same set of epitopes then requires a higher effective resolution; equivalently, a reduced labelling efficiency must be compensated either by higher resolution that can disambiguate sparser localisations or by averaging over more particles. The parameter sweep framework (Fig. 3) is designed to expose these trade-offs, letting the user identify, for a given target feature, the joint operating envelope of probe, labelling, and acquisition parameters within which the feature remains resolvable.

---

**Sup. Fig. S1.**
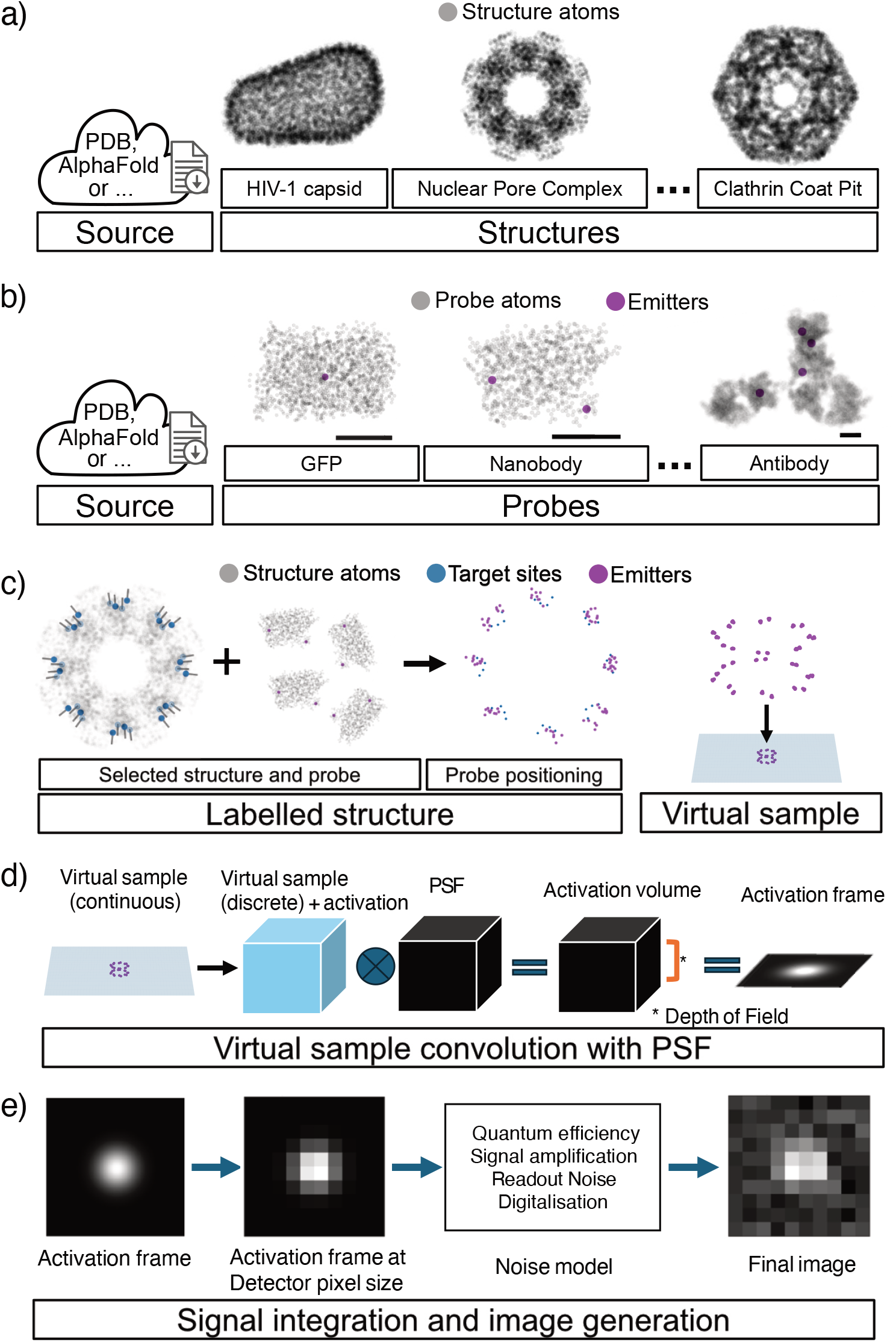
Computational pipeline for image formation simulation. a) Structure selection. VLab4Mic workflow starts by selecting a structure of interest. Here a model for the macromolecular complex is based on PDB/CIF atomic models from available databases such as the PDB or AlphaFold. **b) Probe specification**. We specify a probe model, such as a GFP or an antibody that is conjugated with fluorophores (magenta dots) and binds to specific sites in the structure, known as target sites (not shown). The number of fluorophores shown in the antibody and nanobody models correspond to their degree of labelling. Scalebars = 2 nm. **c) Labelled structure and virtual sample**. Having selected a structure and a probe, VLab4Mic models the labelling of the targets in the structure by positioning copies of the probe onto the target sites while following the geometrical constraints of both models. This creates a general model for this particular labelling strategy. To construct the virtual sample, one or more copies of the labelled structure are placed within a defined volume. Here, each labelled particle is independent of each other. **d) Forward model and activation**. VLab4Mic uses a forward model for image generation to create an imaging simulation of the virtual sample. First, the virtual sample is discretised into a grid whose voxels are of the same dimensions as a 3D PSF model. An activation model assigns a random number of photons to each emitter in the virtual sample, so that the intensity per voxel reflects the total amount of photons emitted in that volume. The discretised virtual sample is then convolved with the 3D PSF to generate an activation volume. In a final step, an activation frame is generated by integrating the signal of the activation volume over the depth of field specified by the imaging modality. **e) Detection and noise**. The activation frame represents the spatial distribution of photons in the field of view that reach the detector. This image is binned (with addition) so that the new pixel size matches the detector pixel size, which now represents the total photon intensity at detection. A modality-dependent noise model is applied to each pixel which includes efficiency of detection, signal amplification, readout and digitalisation noise.

**Sup. Fig. S2.**
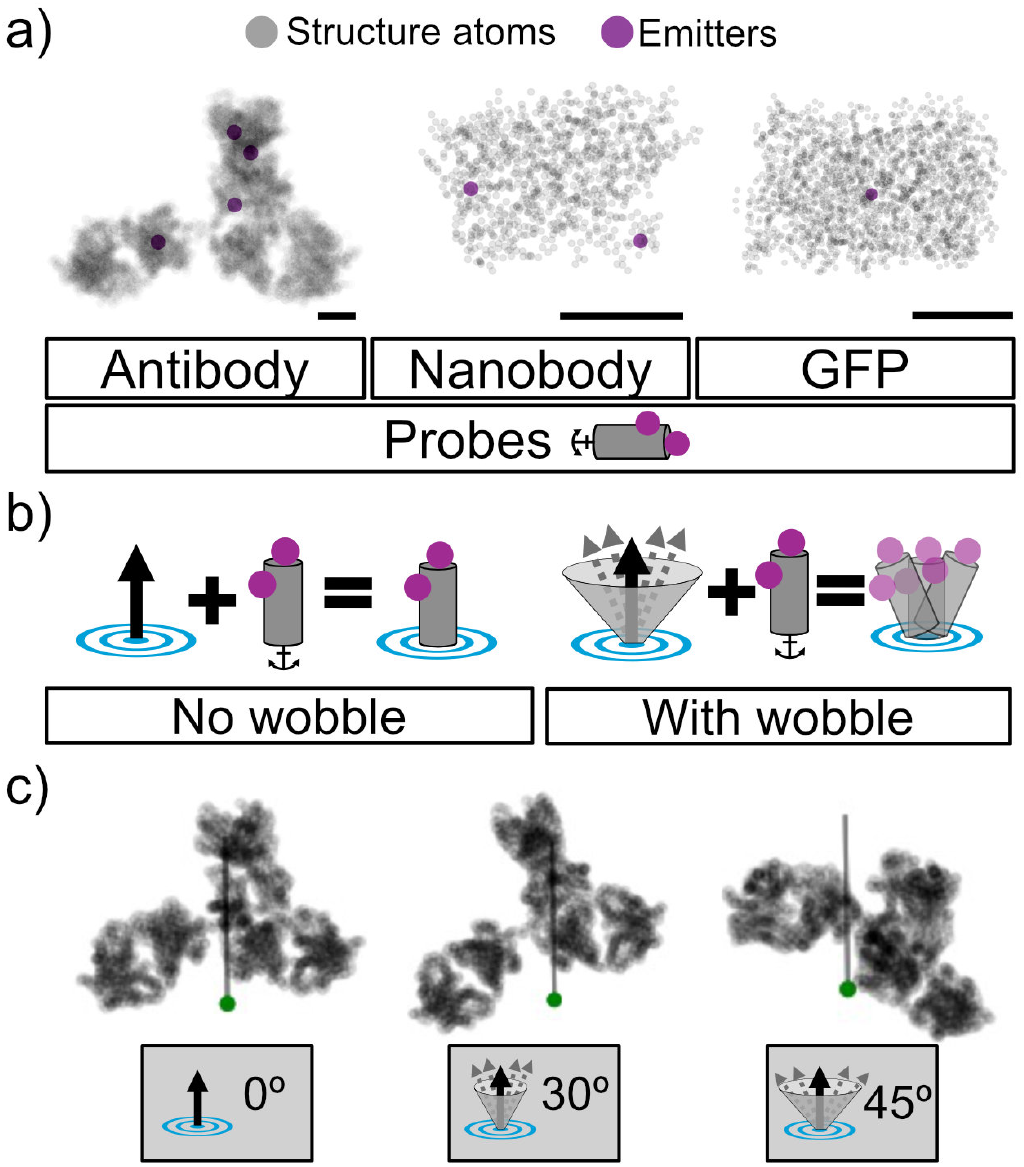
Library of molecular probe models and positioning logic. a) Probe model. A probe is a model for the spatial distribution of emitters or fluorophores (magenta), relative to the probe’s own model (grey). The probe model contains information of a target site for which it has affinity to bind (represented as an anchor at the bottom). This probe is considered a rigid body that can be manipulated based on the anchor and an internal axis. VLab4Mic can model conventional labelling strategies such as antibodies by using the antibody PDB to define putative emitter sites, such as Lysine residues used for NHS-ester fluorophore conjugation. The number of fluorophores shown in the antibody and nanobody models correspond to their degree of labelling. The anchor point can be approximated by known paratope sequences or with a user-defined site. **b) Probe positioning**. We model a probe binding its epitope by repositioning the probe as a rigid body to (1) match the probe anchor and a target site, and (2) align the probe internal axis with the target normal vector. This process can include a wobbling step (right), in which the target normal is given a random perturbation within a user-defined cone as depicted. **c) Antibody probe example**. Model for an antibody probe, depicted from its structural atoms (grey). Its target site (green) and target normal (vertical grey line) are shown to illustrate probe positioning scenarios with no wobble (0 degrees, left), wobble within 30 (centre) or 45 degrees from the target normal. Scale bars in a) are 2 nm.

**Sup. Fig. S3.**
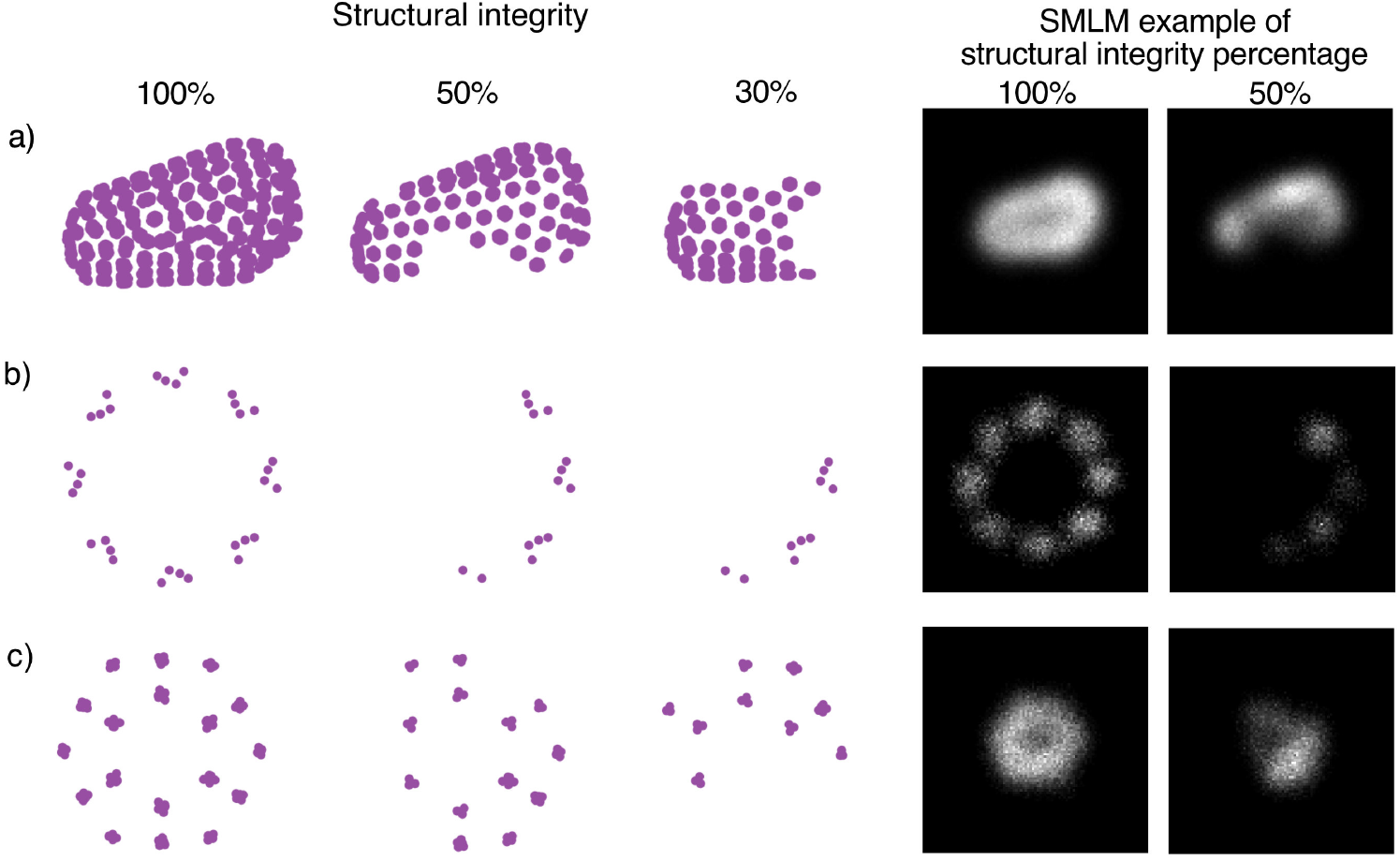
Structural integrity modelling. VLab4Mic can emulate structural integrity as a proxy for epitope accessibility. For each structure, the target positions were used as emitter positions. An example of a labelled particle is shown for different degrees of structural integrity (left panels). SMLM imaging simulations per particle and structural integrity are shown on the right. **a) HIV-1 capsid**. Target sequence: SPRTLNA; short-range cluster = 20 Å, long-range clusters = 100 Å. **b) Nuclear Pore Complex**. Target sequence: ELAVGSL; short-range cluster = 300 Å, long-range clusters = 600 Å. **c) Clathrin Coated Pit**. Target sequence: EQATETQ; short-range cluster = 100 Å, long-range clusters = 200 Å.

**Sup. Fig. S4.**
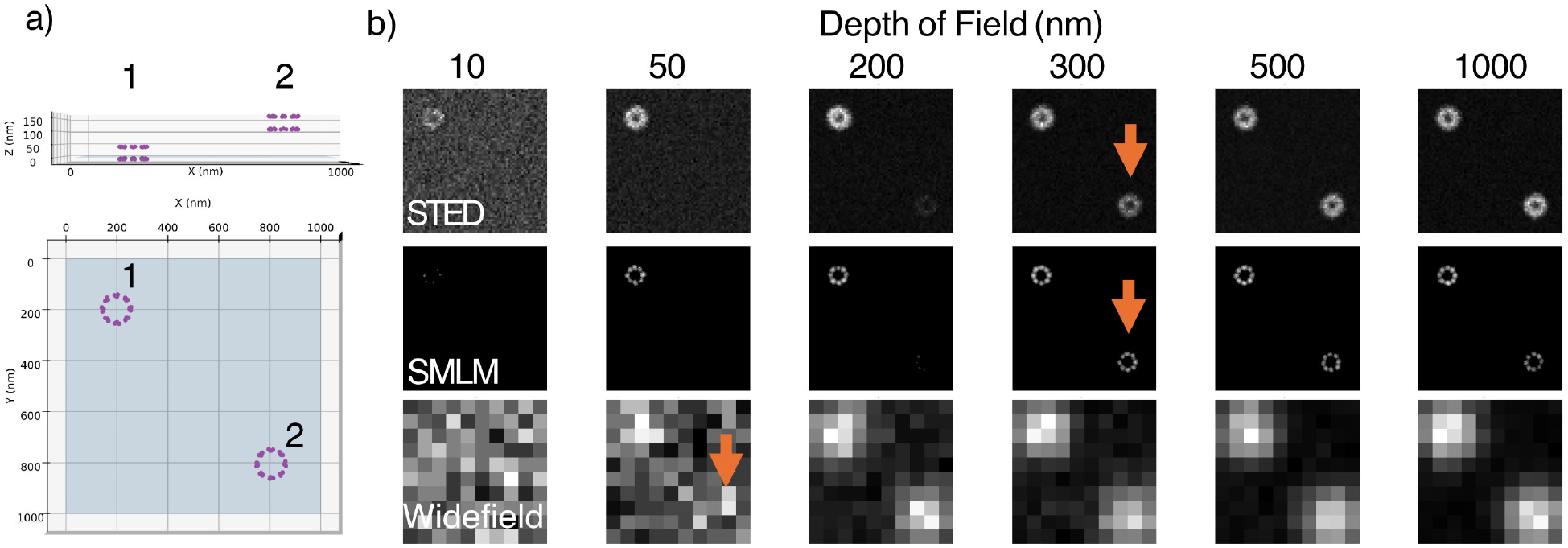
Impact of depth of field on 3D imaging outcomes. a) Virtual sample. Two nuclear pore complexes: NPC #1 located at the focus plane, NPC #2 placed 150 nm above the Z axis and opposite in the FOV. The virtual sample is shown from lateral and top perspectives. **b) Simulated images per modality**. Imaging simulations per modality where the only parameter changed was the depth of field. An orange arrow indicates the depth of field at which a given modality collects signal from NPC #2.

**Sup. Fig. S5.**
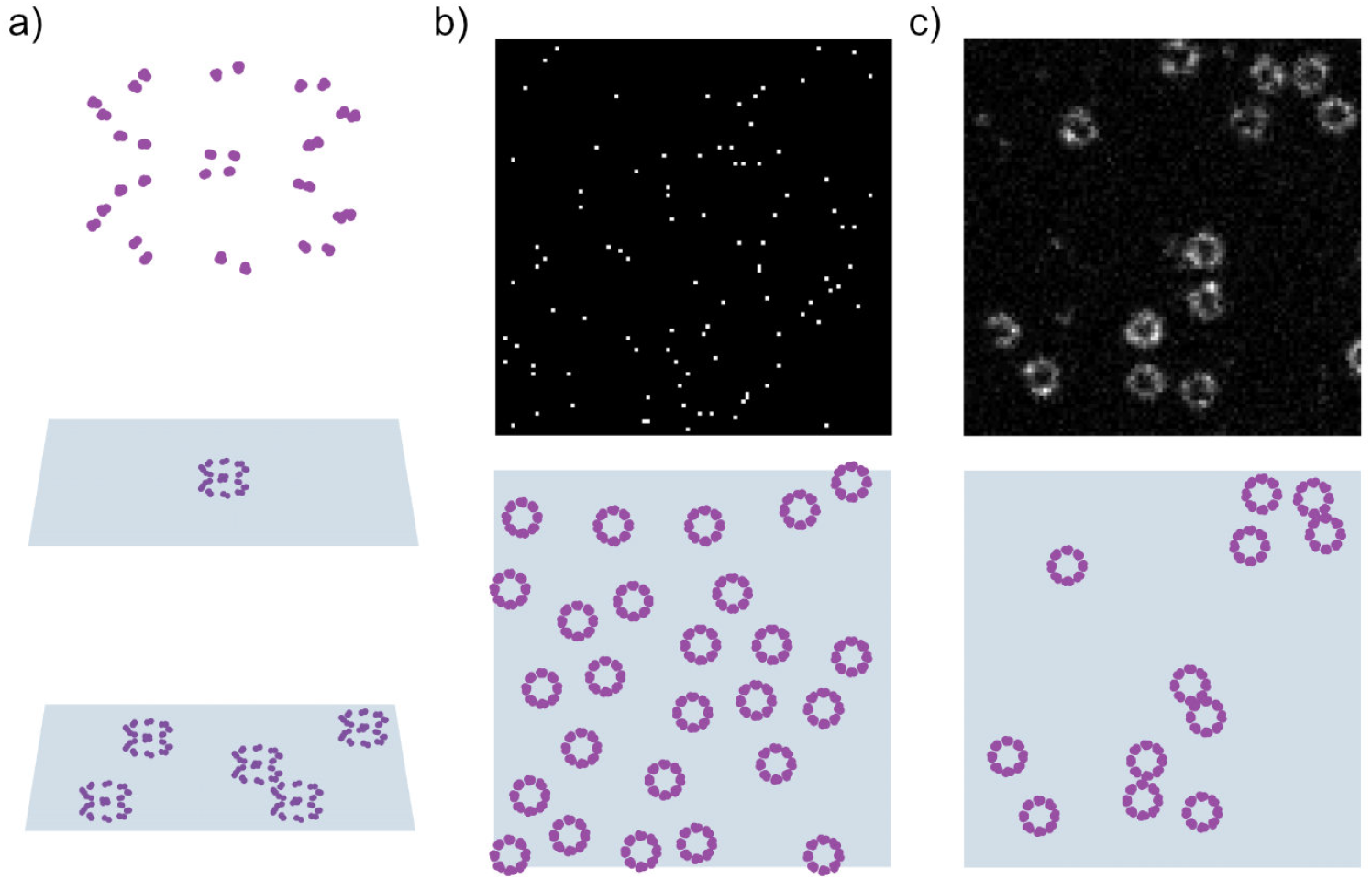
Construction and parameterisation of the 3D virtual sample. a) Sample space and placement. A virtual sample represents fluorophores in a 3D sample space (XY plane highlighted in blue). A labelled structure (top) can be placed within the sample space by specifying relative positions within the field. The default placement is a single labelled structure at the middle of the sample (middle), whereas setting multiple copies generates randomised, non-overlapping positions (bottom). **b) Placement from a binary mask**. Labelled structures can be sampled from a binary mask, where each pixel represents equal probability for a position to occur within that area. **c) Placement from a greyscale image**. Labelled structures can be placed by locating local maxima positions in a greyscale image. This approach approximates the input image with modelled structures.

**Sup. Fig. S6.**
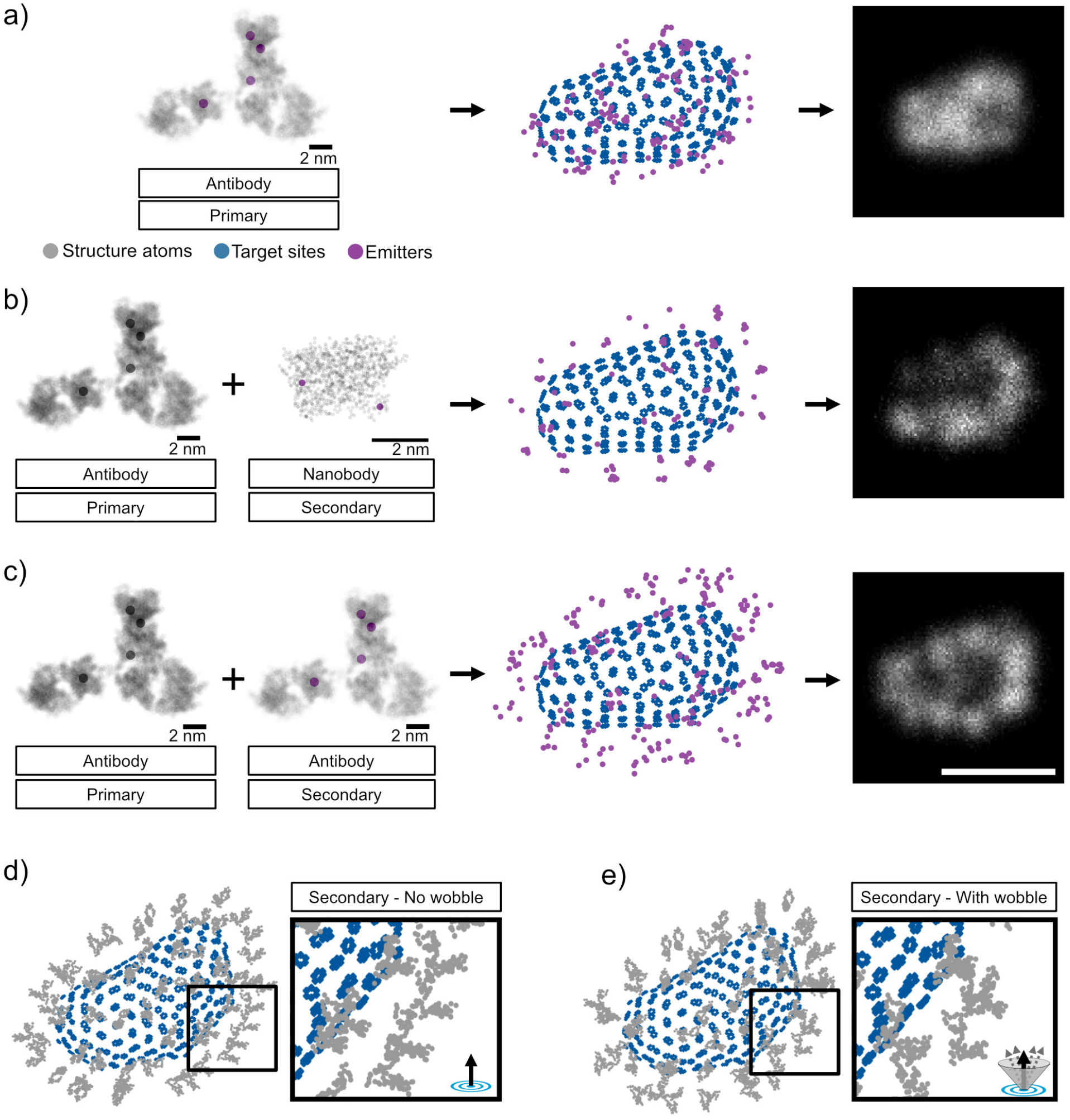
Simulation of primary and secondary linkage error. a) Primary antibody labelling. (Left) Primary antibody with conjugation sites used to define the emitters for this probe. (Middle) Model for the HIV-1 capsid labelled with a primary antibody, showing the target sites of the capsid, the emitters. (Right) SMLM image simulation of HIV-1 capsid labelled with a primary antibody. **b) Primary + secondary nanobody**. (Left) Primary antibody without conjugation sites for emitters, paired with a secondary probe based on a Nanobody whose conjugation sites are used as emitter sites. (Middle) Model for the HIV-1 capsid labelled with the primary probe and a secondary directed to the primary, showing the target sites of the capsid, the emitters. (Right) SMLM image simulation of HIV-1 capsid labelled with a primary and a secondary nanobody. **c) Primary + secondary antibody**. (Left) Primary antibody without conjugation sites for emitters, paired with a secondary probe based on the same model for the primary antibody. (Middle) Model for the HIV-1 capsid labelled with the primary probe and a secondary directed to the primary, showing the target sites of the capsid, the emitters, and an example SMLM image. (Right) SMLM image simulation of HIV-1 capsid labelled with a primary and a secondary antibody. Scalebar = 100 nm. **d) Secondary with no wobble**. Secondary probes are independent of primaries, which allows for independent parameterisation such as degree of wobble. Here, the secondary probe model is represented in grey without fluorophores; primary is not shown. If the secondary is set to not include wobble, the orientation of secondaries will reflect the primary and epitope normal vector. This results in neighbouring secondaries to have similar orientations. **e) Secondary with wobble**. Here the same secondary probe includes a wobble of 60º. Secondary probe is represented in grey without fluorophores, primary is not shown. This results in randomised orientations of the secondary probes.

**Sup. Fig. S7.**
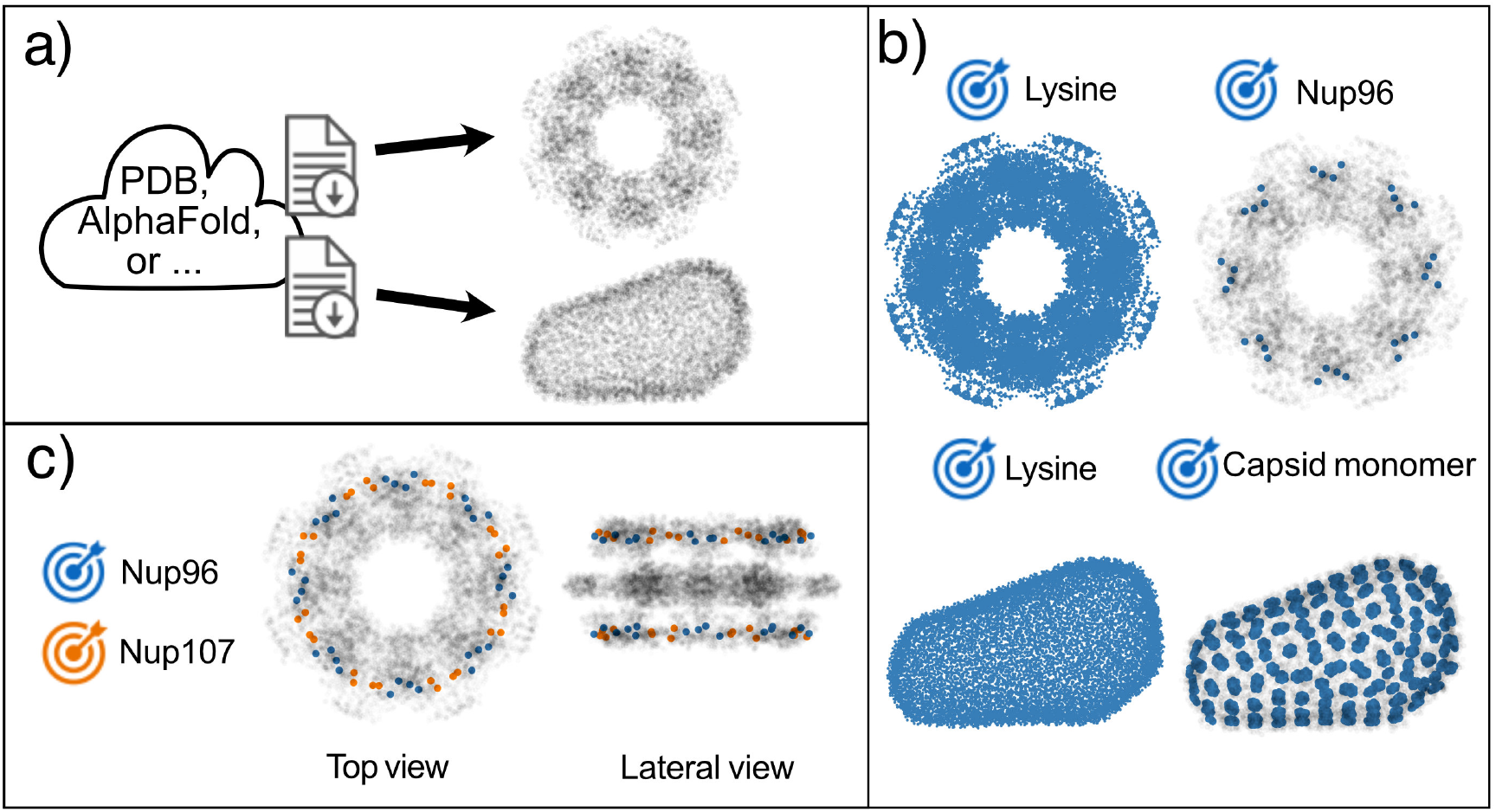
Extraction of ground-truth geometry from atomic PDB models. A structure in VLab4Mic is the representation of an atomic model for the macromolecule of interest. **a) Structure parsing**. VLab4Mic loads PDB files from databases such as the RCSB Protein Data Bank (RCSB PDB) or AlphaFold. We use BioPython parsing tools to read the atoms in the file. For large assemblies, the atoms in the file represent an asymmetric unit (highlighted in blue), as is the case for the Nuclear Pore Complex (NPC) (top). VLab4Mic creates the complete assembly model (grey) from the assembly operations in the structure file. **b) Target parsing**. A VLab4Mic structure provides methods to select positions that match a specific amino-acid sequence or residue. For instance, it is possible to retrieve all Lysine residues found in the NPC structure (top left), or all positions that match the sequence “ELAVGSL”, a peptide motif found in the Nup96 protein. The same procedure can be applied to other structures such as the HIV-1 capsid model (bottom): the example shows the Lysine residues in the HIV-1 capsid on the left, and positions that match the sequence “SPRTLNA”, a peptide found in the HIV-1 capsid monomer (CA). **c) Multiple target selection**. VLab4Mic makes it straightforward to retrieve multiple target sites, each based on its specific property (residues or sequences). Example shows Nup96 and Nup107 C-terminals.

**Sup. Fig. S8.**
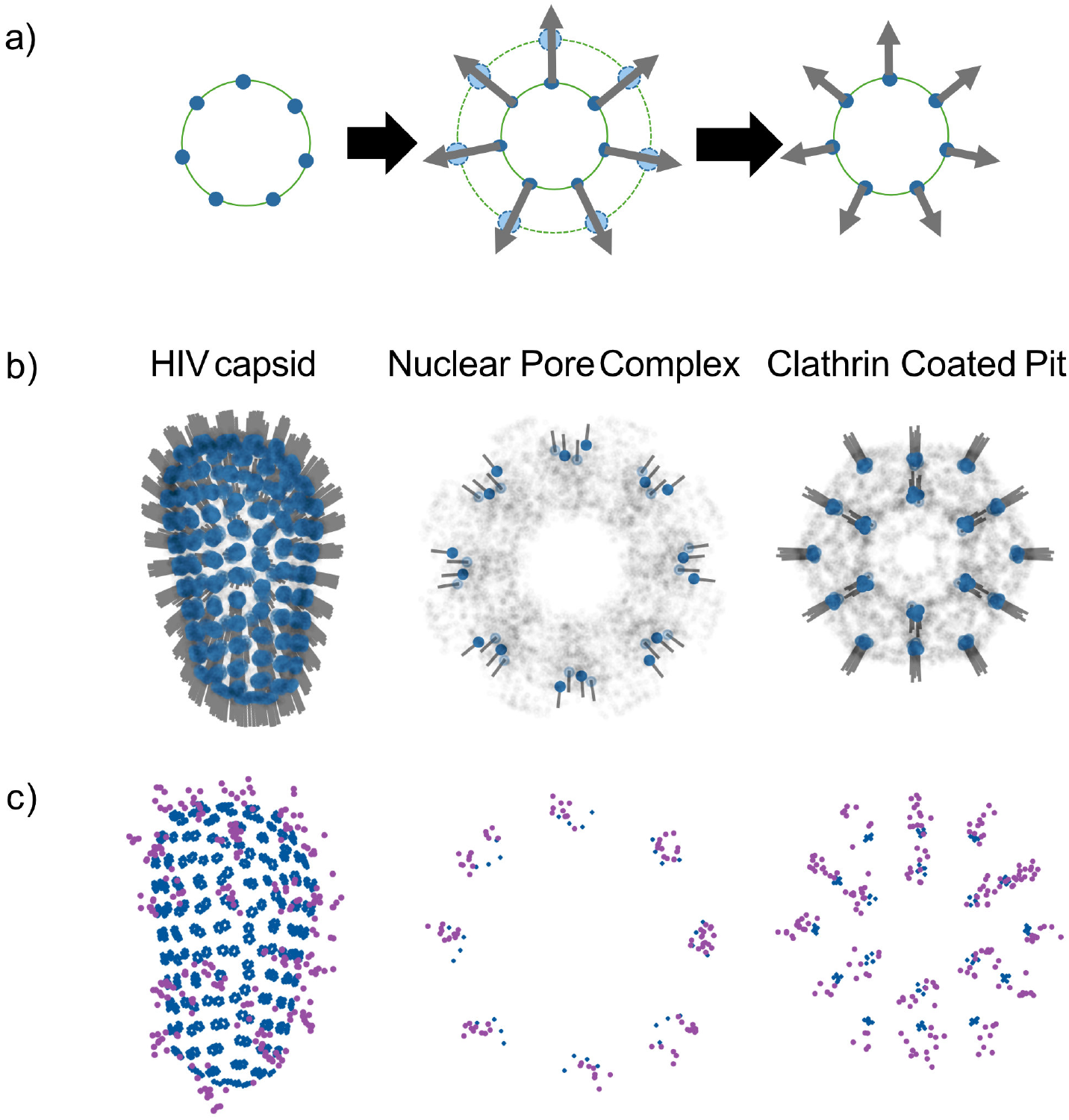
Algorithm for probe positioning and surface orientation. a) Normal vector approximation. We approximate the normal vector direction by creating a scaled-down version of the target sites and defining each normal vector as the vector from each scaled-target back to its corresponding target. **b) Targets and normal vectors across structures**. Each target site (blue) on a structure has its own normal vector approximated as a vector that starts from the target site and points outwards (grey line segment) from the structure. **c) Probe positioning**. Examples of different structures labelled with an antibody probe.

**Sup. Fig. S9.**
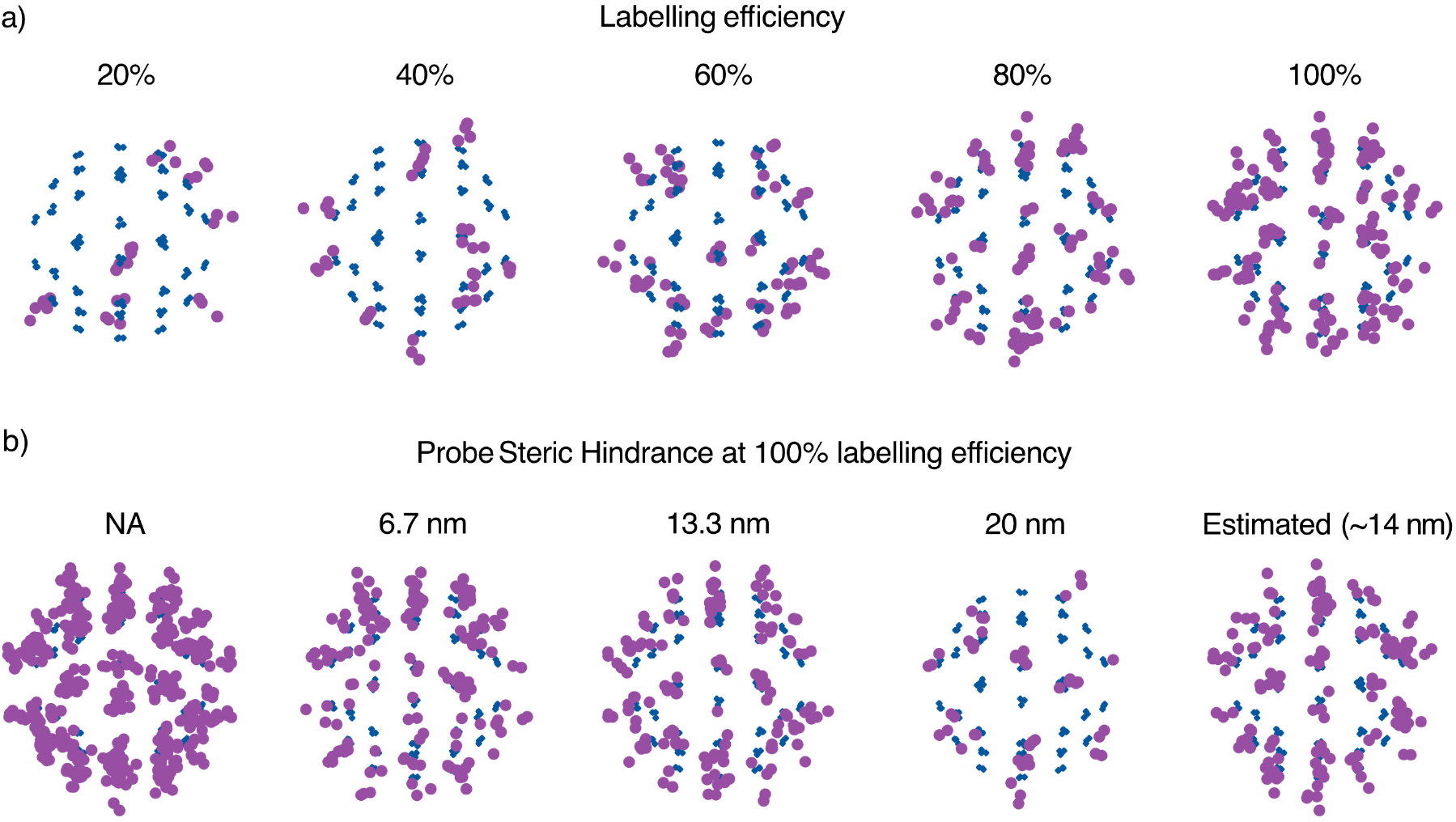
Modelling binding heterogeneity and probe steric hindrance. a) Labelling efficiency. We model labelling efficiency during probe positioning by taking it as the probability of success in a Bernoulli trial per target site. Shown with a linker probe while varying the labelling efficiency from 20% to 100%. In this example, probe binding also takes into account proximity to neighbouring targets. **b) Probe steric hindrance**. We model steric constraints as the minimum distance required for neighbouring epitopes to be available for labelling. For each probe, this distance is calculated from the maximum pairwise distance between its emitters. Shown are examples using no hindrance, different fixed values for hindrance, and the estimated distance from the probe model itself (this calculation is used by default).

**Sup. Fig. S10.**
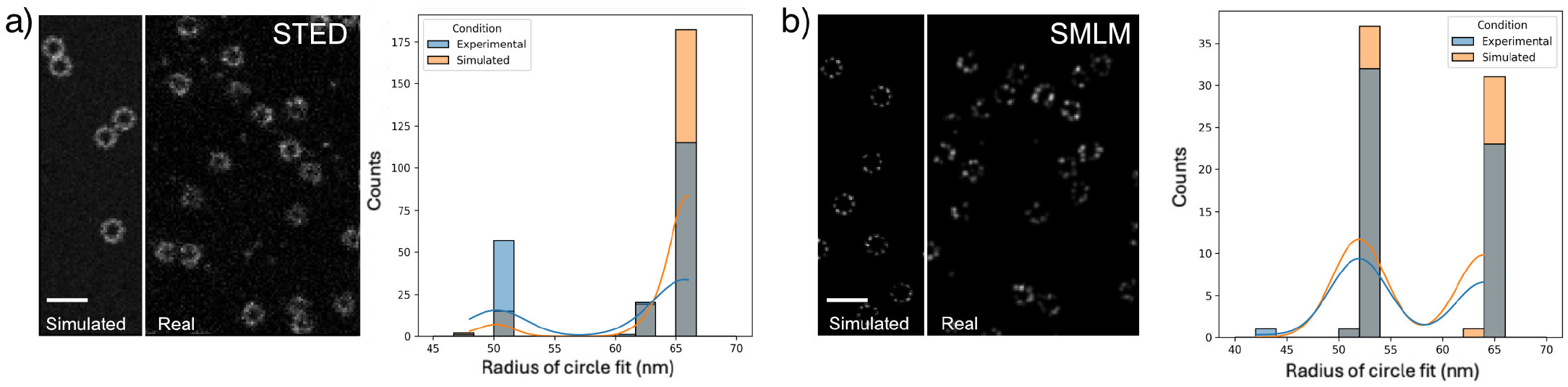
Distribution of radii fit on simulated and experimental images. Image overlays show simulated and experimental data available (36) for **a) STED** or **b) SMLM**, together with their histogram of fitted radii. Parameters were the same per modality. Scale bars: 250 nm.

**Sup. Fig. S11.**
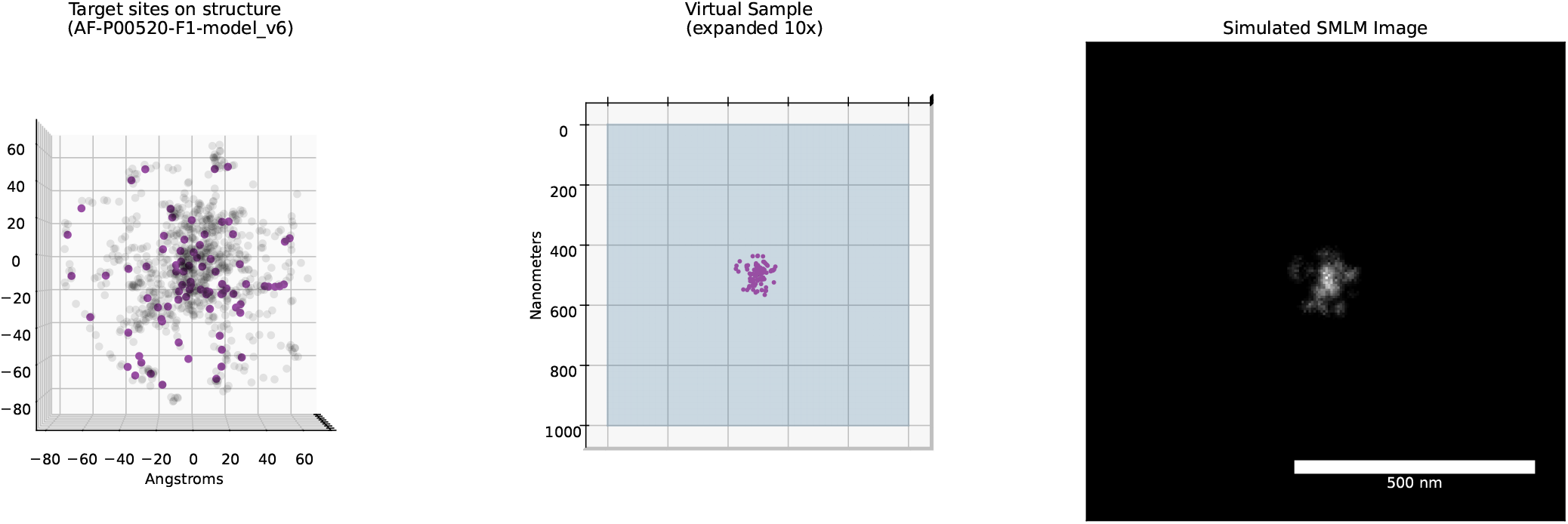
Simulations of AlphaFold models. VLab4Mic can use AlphaFold models as structural inputs for simulations. Here we show an example of an AlphaFold model for the Striatin-interacting protein 1. **Left:** target sites for Lysine residues (magenta) and atoms from the structure (grey). **Middle:** virtual sample with a single copy of the labelled structure. The sample was expanded isotropically with a factor of 10 times. **Right:** SMLM imaging simulation of the virtual sample.

**Sup. Fig. S12.**
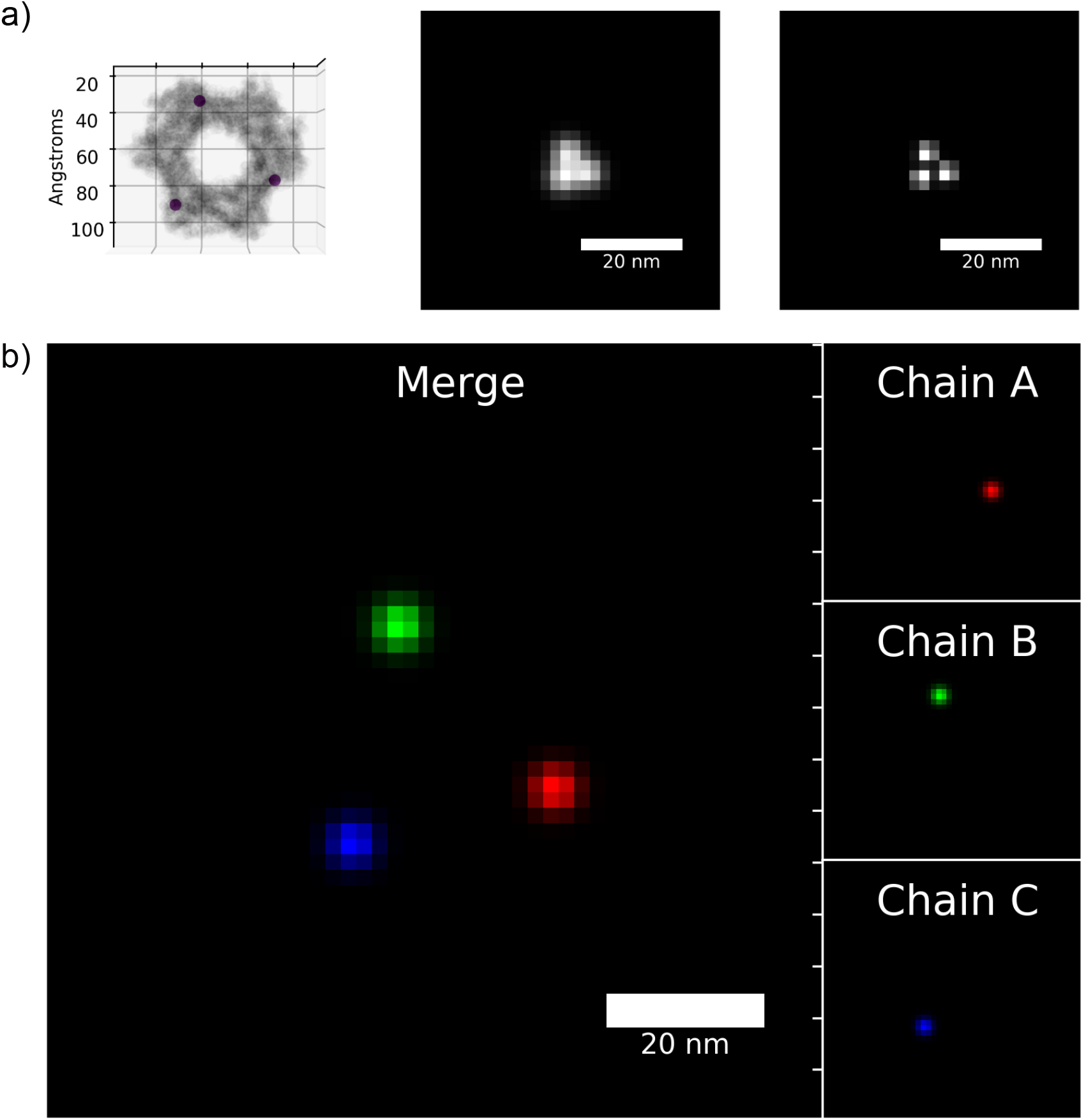
Site-specific labelling and multicolour imaging. VLab4Mic allows users to define specific residue numbers and chains to be targeted within a protein complex. **a) Single-channel site-specific SMLM**. Human homotrimeric proliferating cell nuclear antigen (PCNA) labelled with a probe targeting SER-186 in each chain of the trimer. The leftmost panel shows the target sites for SER-186 residues (magenta) and atoms from the structure (grey). Using these target sites, we show example SMLM imaging simulations of the site-specific labels with resolution of 2 nm and 1 nm. **b) Multicolour site-specific SMLM**. Three different probes are specified for multicolour imaging: each was parameterised to target a different chain of the PCNA trimer at residue SER-186. This results in each chain being imaged in a different channel. For visualisation purposes, the virtual sample was expanded isotropically with a factor of 5. The 3-colour SMLM imaging simulation is shown both as independent channels (each one corresponding to a different chain in the homotrimeric antigen) and as a composite image.

**Sup. Fig. S13.**
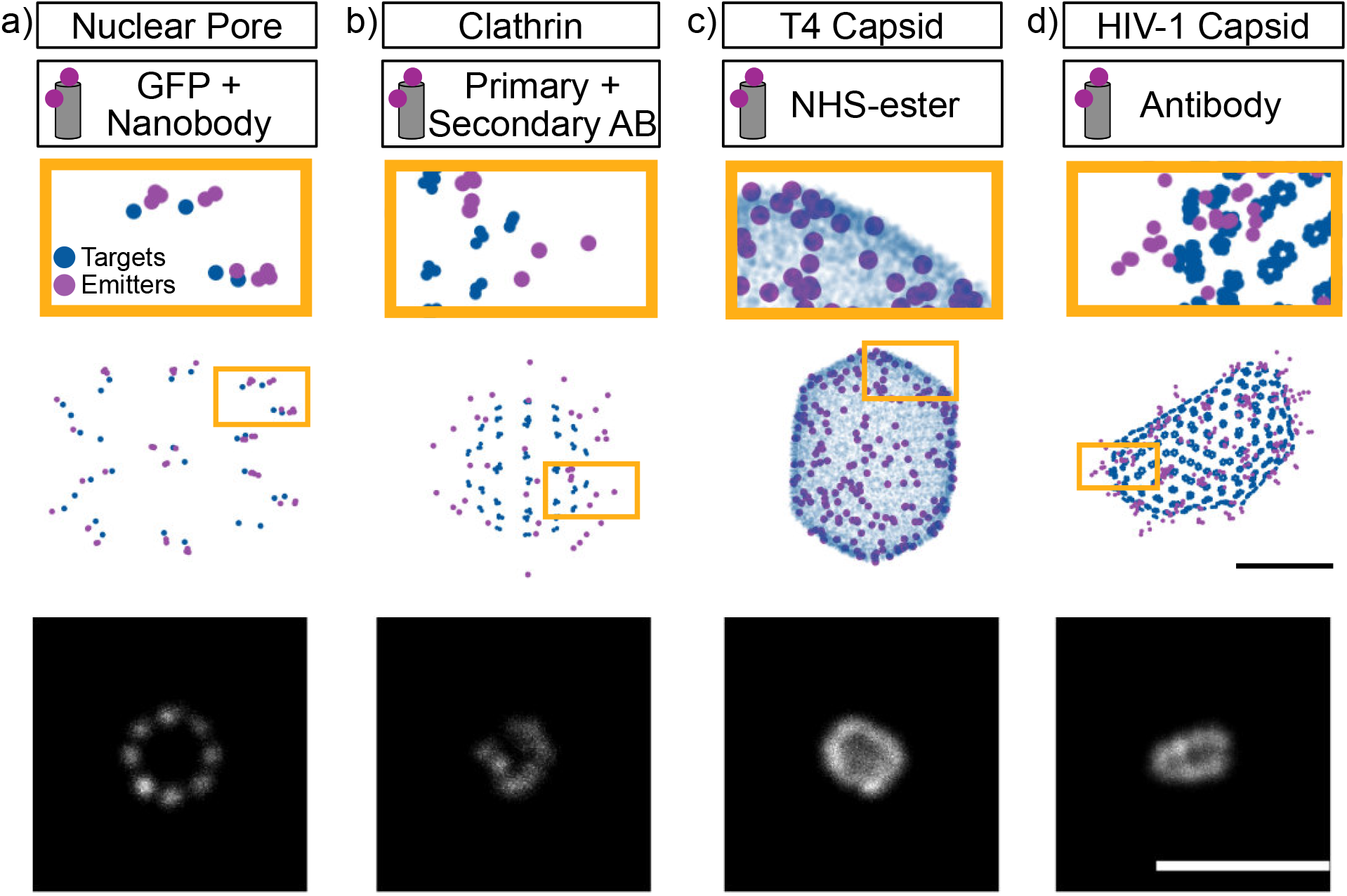
Diverse biological structures and labelling strategies. Four representative examples demonstrate versatility. **a) Nuclear pore complex (7R5K)**. GFP-nanobodies targeting Nup96. **b) Clathrin-coated pit (1XI5)**. Primary-secondary antibody pair targeting clathrin heavy chains. **c) T4 bacteriophage capsid (8GMO)**. NHS-ester conjugation. **d) HIV-1 capsid (3J3Y)**. Antibody labelling of p24. These examples encompass diverse structural scales (from 40 nm clathrin cages to 100 nm phage capsids) and geometries: lattices (clathrin), icosahedral symmetry (T4 phage), conical fullerene capsid (HIV-1), and symmetric ring structures (nuclear pores). Each labelling strategy introduces characteristic fluorophore displacement patterns. GFP-nanobodies provide tight labelling, while primary-secondary antibody pairs introduce larger displacements that affect apparent structure size in super-resolution images. Scalebar on labelled structures is 50 nm; scalebar for image simulation is 250 nm.

**Sup. Video. 1.**
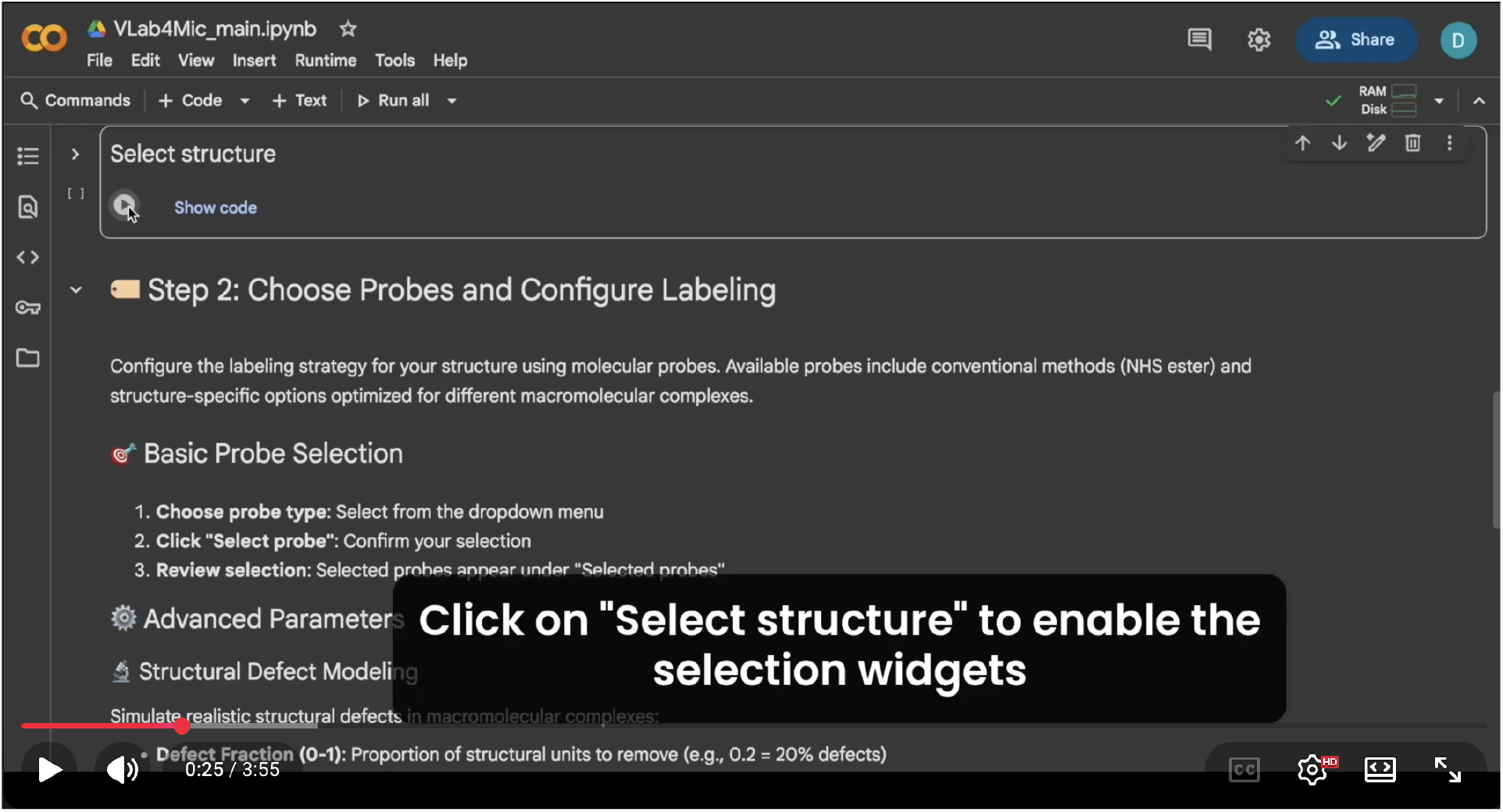
Complete workflow of a virtual microscopy experiment with VLab4Mic. (▶ Watch video) Example use of a VLab4Mic experiment for selection of structure, probe, virtual sample parameters, imaging modalities and acquisition parameters to generate multimodal image simulations.

**Sup. Video. 2.**
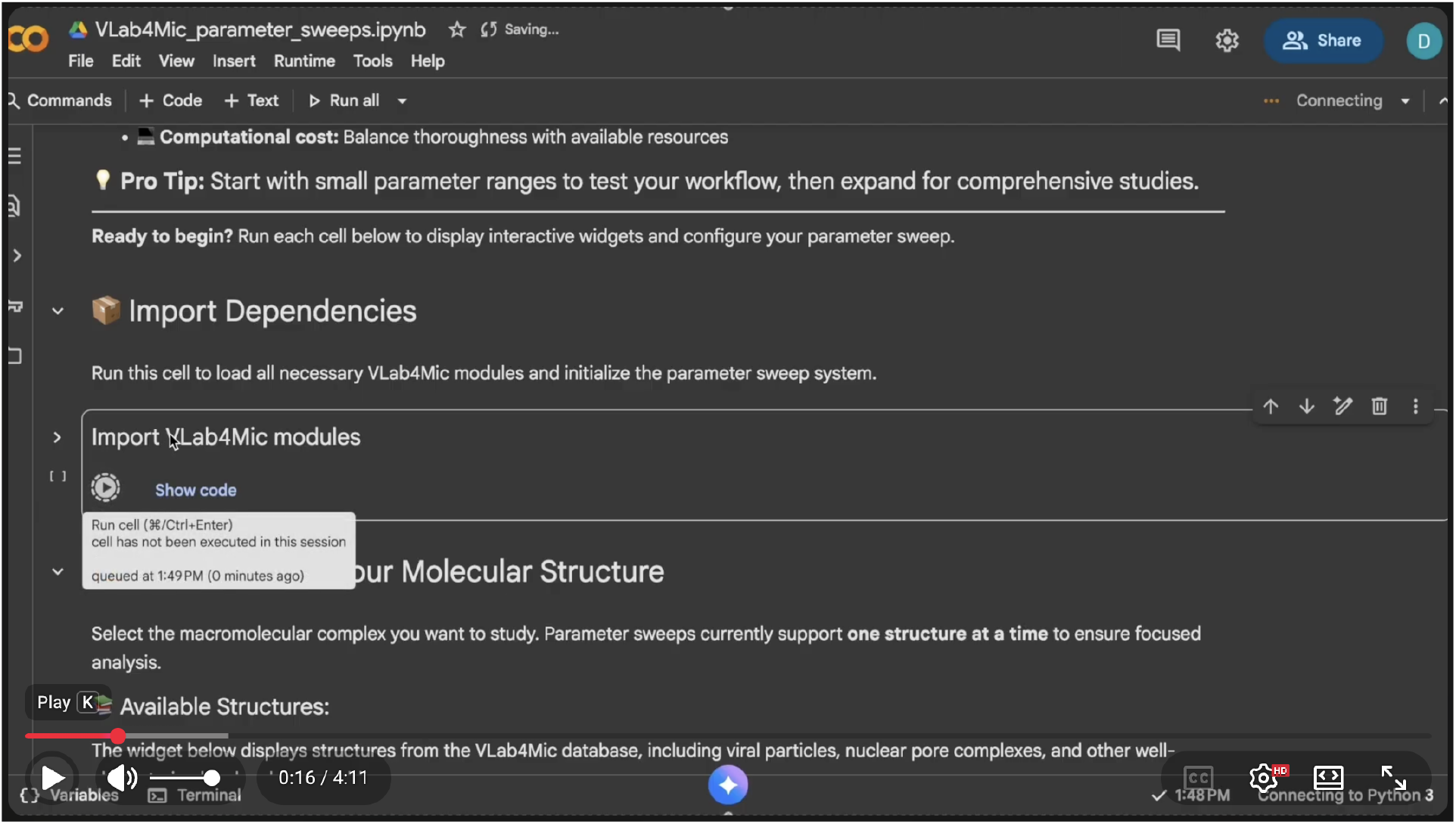
Parameter sweep analysis with codeless notebooks. (▶ Watch video) Example use of the VLab4Mic parameter sweep functionality to generate extensive simulation datasets from parameter combinations. The resulting dataset is analysed by comparing each parameter combination output against a reference standard.

**Sup. Video.3.**
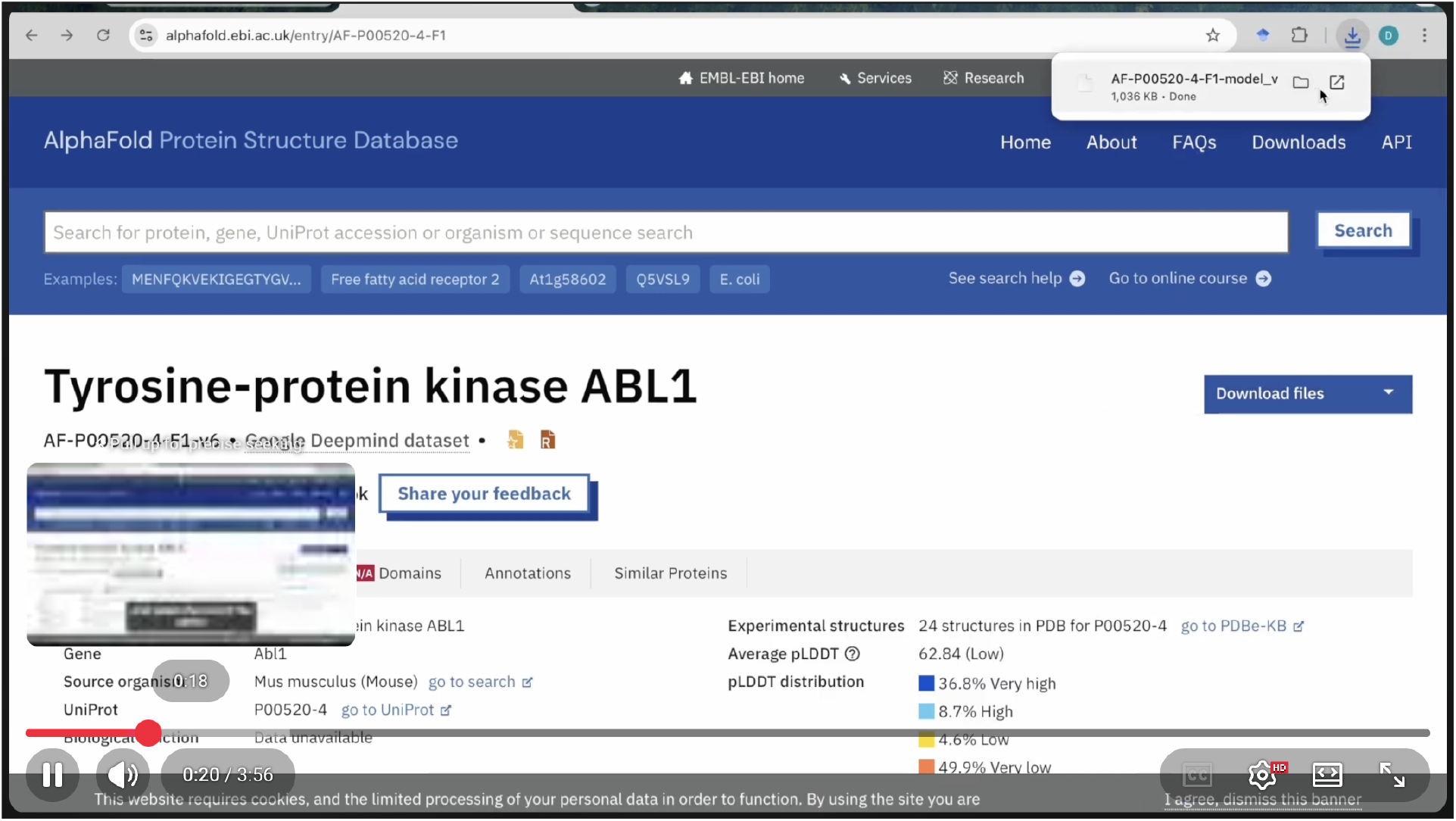
Use of AlphaFold-predicted models in VLab4Mic. (▶ Watch video) A model predicted by AlphaFold can be used as an input structure for VLab4Mic modelling and simulations. Models from other databases or custom models made by users can also be used as input structures.

## Bibliography

1. Jervis Vermal Thevathasan, Maurice Kahnwald, Konstanty Cieśliński, Philipp Hoess, Sudheer Kumar Peneti, Manuel Reitberger, Daniel Heid, Krishna Chaitanya Kasuba, Sarah Janice Hoerner, Yiming Li, Yu-Le Wu, Markus Mund, Ulf Matti, Pedro Matos Pereira, Ricardo Henriques, Bianca Nijmeijer, Moritz Kueblbeck, Vilma Jimenez Sabinina, Jan Ellenberg, and Jonas Ries. Nuclear pores as versatile reference standards for quantitative superresolution microscopy. Nature Methods, 16(10):1045–1053, October 2019. ISSN 1548-7105. doi:10.1038/s41592-019-0574-9.

2. Lothar Schermelleh, Alexia Ferrand, Thomas Huser, Christian Eggeling, Markus Sauer, Oliver Biehlmaier, and Gregor P. C. Drummen. Super-resolution microscopy demystified. Nature Cell Biology, 21(1):72–84, January 2019. ISSN 1476-4679. doi:10.1038/s41556-018-0251-8.

3. Mickaël Lelek, Melina T. Gyparaki, Gerti Beliu, Florian Schueder, Juliette Griffié, Suliana Manley, Ralf Jungmann, Markus Sauer, Melike Lakadamyali, and Christophe Zimmer. Single-molecule localization microscopy. Nature Reviews Methods Primers, 1(1):39, June 2021. ISSN 2662-8449. doi:10.1038/s43586-021-00038-x.

4. Dominique Bourgeois. Single molecule imaging simulations with advanced fluorophore photophysics. Communications Biology, 6(1):53, January 2023. ISSN 2399-3642. doi:10.1038/s42003-023-04432-x.

5. J. Griffié, T. A. Pham, C. Sieben, R. Lang, V. Cevher, S. Holden, M. Unser, S. Manley, and D. Sage. Virtual-SMLM, a virtual environment for real-time interactive SMLM acquisition, March 2020. doi:10.1101/2020.03.05.967893.

6. Tibor Novák, Tamás Gajdos, József Sinkó, Gábor Szabó, and Miklós Erdélyi. TestSTORM: Versatile simulator software for multimodal super-resolution localization fluorescence microscopy. Scientific Reports, 7(1):951, April 2017. ISSN 2045-2322. doi:10.1038/s41598-017-01122-7.

7. Varun Venkataramani, Frank Herrmannsdörfer, Mike Heilemann, and Thomas Kuner. SuReSim: Simulating localization microscopy experiments from ground truth models. Nature Methods, 13(4):319–321, April 2016. ISSN 1548-7105. doi:10.1038/nmeth.3775.

8. Jonas Ries. SMAP: A modular super-resolution microscopy analysis platform for SMLM data. Nature Methods, 17(9):870–872, September 2020. ISSN 1548-7105. doi:10.1038/s41592-020-0938-1.

9. Daniel Sage, Thanh-An Pham, Hazen Babcock, Tomas Lukes, Thomas Pengo, Jerry Chao, Ramraj Velmurugan, Alex Herbert, Anurag Agrawal, Silvia Colabrese, Ann Wheeler, Anna Archetti, Bernd Rieger, Raimund Ober, Guy M. Hagen, Jean-Baptiste Sibarita, Jonas Ries, Ricardo Henriques, Michael Unser, and Seamus Holden. Super-resolution fight club: Assessment of 2D and 3D single-molecule localization microscopy software. Nature Methods, 16(5):387–395, May 2019. ISSN 1548-7105. doi:10.1038/s41592-019-0364-4.

10. Manya Kapoor and Mustafa Mir. Simulation of transcription factor clustering in nuclei from molecular kinetics. bioRxiv, 2026. doi:10.64898/2026.06.04.730171. Preprint.

11. Susanna M. Früh, Ulf Matti, Philipp R. Spycher, Marina Rubini, Sebastian Lickert, Thomas Schlichthaerle, Ralf Jungmann, Viola Vogel, Jonas Ries, and Ingmar Schoen. Site-Specifically-Labeled Antibodies for Super-Resolution Microscopy Reveal In Situ Linkage Errors. ACS Nano, 15(7):12161–12170, July 2021. ISSN 1936-0851. doi:10.1021/acsnano.1c03677.

12. Maria Theiss, Jean-Karim Hériché, Craig Russell, David Helekal, Alisdair Soppitt, Jonas Ries, Jan Ellenberg, Alvis Brazma, and Virginie Uhlmann. Simulating structurally variable nuclear pore complexes for microscopy. Bioinformatics, 39(10):btad587, October 2023. ISSN 1367-4811. doi:10.1093/bioinformatics/btad587.

13. Stefan W. Hell and Jan Wichmann. Breaking the diffraction resolution limit by stimulated emission: Stimulated-emission-depletion fluorescence microscopy. Optics Letters, 19(11): 780–782, June 1994. ISSN 1539-4794. doi:10.1364/OL.19.000780.

14. Eric Betzig, George H. Patterson, Rachid Sougrat, O. Wolf Lindwasser, Scott Olenych, Juan S. Bonifacino, Michael W. Davidson, Jennifer Lippincott-Schwartz, and Harald F. Hess. Imaging Intracellular Fluorescent Proteins at Nanometer Resolution. Science, 313(5793): 1642–1645, September 2006. doi:10.1126/science.1127344.

15. Michael J. Rust, Mark Bates, and Xiaowei Zhuang. Sub-diffraction-limit imaging by stochastic optical reconstruction microscopy (STORM). Nature Methods, 3(10):793–796, October 2006. ISSN 1548-7105. doi:10.1038/nmeth929.

16. Samuel T. Hess, Thanu P. K. Girirajan, and Michael D. Mason. Ultra-High Resolution Imaging by Fluorescence Photoactivation Localization Microscopy. Biophysical Journal, 91(11):4258– 4272, December 2006. ISSN 0006-3495, 1542-0086. doi:10.1529/biophysj.106.091116.

17. Helen M. Berman, John Westbrook, Zukang Feng, Gary Gilliland, T. N. Bhat, Helge Weissig, Ilya N. Shindyalov, and Philip E. Bourne. The Protein Data Bank. Nucleic Acids Research, 28(1):235–242, January 2000. ISSN 0305-1048. doi:10.1093/nar/28.1.235.

18. Kazuki Obashi, Kem A. Sochacki, Marie-Paule Strub, and Justin W. Taraska. A conformational switch in clathrin light chain regulates lattice structure and endocytosis at the plasma membrane of mammalian cells. Nature Communications, 14(1):732, February 2023. ISSN 2041-1723. doi:10.1038/s41467-023-36304-7.

19. Thomas D. Goddard, Conrad C. Huang, Elaine C. Meng, Eric F. Pettersen, Gregory S. Couch, John H. Morris, and Thomas E. Ferrin. UCSF ChimeraX: Meeting modern challenges in visualization and analysis. Protein Science: A Publication of the Protein Society, 27(1): 14–25, January 2018. ISSN 1469-896X. doi:10.1002/pro.3235.

20. Kem A. Sochacki, Bridgette L. Heine, Gideon J. Haber, John R. Jimah, Bijeta Prasai, Marco A. Alfonzo-Méndez, Aleah D. Roberts, Agila Somasundaram, Jenny E. Hinshaw, and Justin W. Taraska. The structure and spontaneous curvature of clathrin lattices at the plasma membrane. Developmental Cell, 56(8):1131–1146.e3, April 2021. doi:10.1016/j.devcel.2021.03.017.

21. John S. H. Danial. Super-resolution microscopy for structural biology. Nature Methods, 22 (8):1636–1652, August 2025. ISSN 1548-7105. doi:10.1038/s41592-025-02731-1.

22. Siân Culley, David Albrecht, Caron Jacobs, Pedro Matos Pereira, Christophe Leterrier, Jason Mercer, and Ricardo Henriques. Quantitative mapping and minimization of super-resolution optical imaging artifacts. Nature Methods, 15(4):263–266, April 2018. ISSN 1548-7105. doi:10.1038/nmeth.4605.

23. Zhou Wang, A. C. Bovik, H. R. Sheikh, and E. P. Simoncelli. Image quality assessment: From error visibility to structural similarity. IEEE Transactions on Image Processing, 13(4): 600–612, 2004. doi:10.1109/TIP.2003.819861.

24. Surbhi Shirgill, Daniel J. Nieves, Jeremy Pike, Mohamed Ahmed, Helen Abbott, Mohammed Baragilly, Karoll Savoye, Mark C. Wales, Adam Kaminer, Ruby Peters, Ricardo Henriques, Steven F. Lee, Patrick Rubin-Delanchy, and Dylan M. Owen. Nano-org, a functional resource for single-molecule localisation microscopy data. Nature Communications, 16:8674, 2025. doi:10.1038/s41467-025-63674-x.

25. Surbhi Shirgill, L. G. Jensen, Daniel J. Nieves, Mark C. Wales, Adam Kaminer, Karoll Savoye, Ruby Peters, Mike Heilemann, Ricardo Henriques, Steven F. Lee, Patrick Rubin-Delanchy, and Dylan M. Owen. Nanoscale spatial-omics via contrastive embedding of single-molecule localisation data. bioRxiv, 2025. doi:10.1101/2025.10.07.679170. Preprint.

26. Romain F. Laine, Kalina L. Tosheva, Nils Gustafsson, Robert D. M. Gray, Pedro Almada, David Albrecht, Gail Tournaviti Risa, Fredrik Hurtig, Ann-Christin Lindås, Buzz Baum, Jason Mercer, Christophe Leterrier, Pedro M. Pereira, Siân Culley, and Ricardo Henriques. NanoJ: A high-performance open-source super-resolution microscopy toolbox. Journal of Physics D: Applied Physics, 52(16):163001, February 2019. ISSN 0022-3727. doi:10.1088/1361-6463/ab0261.

27. Yu-Le Wu, Philipp Hoess, Aline Tschanz, Ulf Matti, Markus Mund, and Jonas Ries. Maximum-likelihood model fitting for quantitative analysis of SMLM data. Nature Methods, 20(1): 139–148, 2023. doi:10.1038/s41592-022-01676-z.

28. Lucas von Chamier, Romain F. Laine, Johanna Jukkala, Christoph Spahn, Daniel Krentzel, Elias Nehme, Martina Lerche, Sara Hernández-Pérez, Pieta K. Mattila, Eleni Karinou, Seámus Holden, Ahmet Can Solak, Alexander Krull, Tim-Oliver Buchholz, Martin L. Jones, Loic A. Royer, Christophe Leterrier, Yoav Shechtman, Florian Jug, Mike Heilemann, Guillaume Jacquemet, and Ricardo Henriques. Democratising deep learning for microscopy with ZeroCostDL4Mic. Nature Communications, 12(1):2276, April 2021. ISSN 2041-1723. doi:10.1038/s41467-021-22518-0.

29. Artur Speiser, Lucas-Raphael Müller, Philipp Hoess, Ulf Matti, Christopher J. Obara, Wesley R. Legant, Anna Kreshuk, Jakob H. Macke, Jonas Ries, and Srinivas C. Turaga. Deep learning enables fast and dense single-molecule localization with high accuracy. Nature Methods, 18(9):1082–1090, 2021. doi:10.1038/s41592-021-01236-x.

30. Sheng Liu, Philipp Hoess, and Jonas Ries. Super-Resolution Microscopy for Structural Cell Biology. Annual Review of Biophysics, 51(1):301–326, May 2022. ISSN 1936-122X, 1936-1238. doi:10.1146/annurev-biophys-102521-112912.

31. Francisco Balzarotti, Yvan Eilers, Klaus C. Gwosch, Arvid H. Gynnå, Volker Westphal, Fernando D. Stefani, Johan Elf, and Stefan W. Hell. Nanometer resolution imaging and tracking of fluorescent molecules with minimal photon fluxes. Science, 355(6325):606–612, February 2017. doi:10.1126/science.aak9913.

32. Kirti Prakash. At the molecular resolution with MINFLUX? Philosophical Transactions of the Royal Society A: Mathematical, Physical and Engineering Sciences, 380(2220):20200145, February 2022. doi:10.1098/rsta.2020.0145.

33. Susanne C. M. Reinhardt, Luciano A. Masullo, Isabelle Baudrexel, Philipp R. Steen, Rafal Kowalewski, Alexandra S. Eklund, Sebastian Strauss, Eduard M. Unterauer, Thomas Schlichthaerle, Maximilian T. Strauss, Christian Klein, and Ralf Jungmann. ångström-resolution fluorescence microscopy. Nature, 617(7962):711–716, May 2023. ISSN 1476-4687. doi:10.1038/s41586-023-05925-9.

34. Niels Radmacher, Alexey I. Chizhik, Oleksii Nevskyi, José Ignacio Gallea, Ingo Gregor, and Jörg Enderlein. Molecular Level Super-Resolution Fluorescence Imaging. Annual Review of Biophysics, 54(1):163–184, May 2025. ISSN 1936-122X, 1936-1238. doi:10.1146/annurev-biophys-071524-105321.

35. Bruno M. Saraiva, António D. Brito, Guillaume Jacquemet, and Ricardo Henriques. Rxiv-maker: an automated template engine for streamlined scientific publications, 2025. doi:10.48550/arXiv.2508.00836.

36. Jervis Vermal Thevathasan, Maurice Kahnwald, Konstanty Cieśliński, Philipp Hoess, Sudheer Kumar Peneti, Manuel Reitberger, Daniel Heid, Krishna Chaitanya Kasuba, Sarah Janice Hoerner, Yiming Li, Yu-Le Wu, Markus Mund, Ulf Matti, Pedro Matos Pereira, Ricardo Henriques, Bianca Nijmeijer, Moritz Kueblbeck, Vilma Jimenez Sabinina, Jan Ellenberg, and Jonas Ries. Nuclear pores as versatile reference standards for quantitative superresolution microscopy (biostudies dataset deposit). https://www.ebi.ac.uk/biostudies/bioimages/studies/S-BIAD8, 2019. BioStudies accession S-BIAD8.

37. Peter J. A. Cock, Tiago Antao, Jeffrey T. Chang, Brad A. Chapman, Cymon J. Cox, Andrew Dalke, Iddo Friedberg, Thomas Hamelryck, Frank Kauff, Bartek Wilczynski, and Michiel J. L. de Hoon. Biopython: Freely available Python tools for computational molecular biology and bioinformatics. Bioinformatics, 25(11):1422–1423, June 2009. ISSN 1367-4803. doi:10.1093/bioinformatics/btp163.

38. Iván Hidalgo-Cenalmor, Marcela Xiomara Rivera Pineda, Bruno M. Saraiva, Ricardo Henriques, and Guillaume Jacquemet. Packaging jupyter notebooks as installable desktop apps using labconstrictor, 2026. doi:10.48550/arXiv.2603.10704.

39. Bruno M. Saraiva, Iván Hidalgo-Cenalmor, António D. Brito, Damián Martínez, Tayla Shakespeare, Guillaume Jacquemet, and Ricardo Henriques. EZInput: A Cross-Environment Python Library for Easy UI Generation in Scientific Computing, January 2026. doi:10.48550/arXiv.2601.08859.

40. Martin Ester, Hans-Peter Kriegel, Jörg Sander, and Xiaowei Xu. A density-based algorithm for discovering clusters in large spatial databases with noise. In Proceedings of the Second International Conference on Knowledge Discovery and Data Mining (KDD-96), pages 226– 231, 1996. doi:10.5555/3001460.3001507.

41. Mohamadreza Fazel, Kristin S. Grussmayer, Boris Ferdman, Aleksandra Radenovic, Yoav Shechtman, Jörg Enderlein, and Steve Pressé. Fluorescence microscopy: A statistics-optics perspective. Reviews of Modern Physics, 96(2):025003, June 2024. doi:10.1103/RevModPhys.96.025003.

42. Bo Zhang, Josiane Zerubia, and Jean-Christophe Olivo-Marin. Gaussian approximations of fluorescence microscope point-spread function models. Applied Optics, 46(10):1819–1829, 2007. doi:10.1364/AO.46.001819.

43. Shyamal Mosalaganti, Agnieszka Obarska-Kosinska, Marc Siggel, Reiya Taniguchi, Beata Turoňová, Christian E. Zimmerli, Katarzyna Buczak, Florian H. Schmidt, Erica Margiotta, Marie-Therese Mackmull, Wim J. H. Hagen, Gerhard Hummer, Jan Kosinski, and Martin Beck. I-based structure prediction empowers integrative structural analysis of human nuclear pores. Science, 376(6598):eabm9506, June 2022. doi:10.1126/science.abm9506.

44. Alexander Fotin, Yifan Cheng, Nikolaus Grigorieff, Thomas Walz, Stephen C. Harrison, and Tomas Kirchhausen. Structure of an auxilin-bound clathrin coat and its implications for the mechanism of uncoating. Nature, 432(7017):649–653, December 2004. ISSN 1476-4687. doi:10.1038/nature03078.

45. Jingen Zhu, Himanshu Batra, Neeti Ananthaswamy, Marthandan Mahalingam, Pan Tao, Xiaorong Wu, Wenzheng Guo, Andrei Fokine, and Venigalla B. Rao. Design of bacteriophage T4-based artificial viral vectors for human genome remodeling. Nature Communications, 14 (1):2928, May 2023. ISSN 2041-1723. doi:10.1038/s41467-023-38364-1.

46. Gongpu Zhao, Juan R. Perilla, Ernest L. Yufenyuy, Xin Meng, Bo Chen, Jiying Ning, Jinwoo Ahn, Angela M. Gronenborn, Klaus Schulten, Christopher Aiken, and Peijun Zhang. Mature HIV-1 capsid structure by cryo-electron microscopy and all-atom molecular dynamics. Nature, 497(7451):643–646, May 2013. ISSN 1476-4687. doi:10.1038/nature12162.

47. Jonas Ries, Christian Kaplan, Evgenia Platonova, Hooman Eghlidi, and Helge Ewers. A simple, versatile method for GFP-based super-resolution microscopy via nanobodies. Nature Methods, 9(6):582–584, June 2012. doi:10.1038/nmeth.1991.

48. Ingmar Schoen, Jonas Ries, Enrico Klotzsch, Helge Ewers, and Viola Vogel. Binding-Activated Localization Microscopy of DNA Structures. Nano Letters, 11(9):4008–4011, September 2011. doi:10.1021/nl2025954.

49. Dominic A. Helmerich, Made Budiarta, Danush Taban, Sören Doose, Gerti Beliu, and Markus Sauer. PCNA as Protein-Based Nanoruler for Sub-10 nm Fluorescence Imaging. Advanced Materials, 36(7):2310104, 2024. ISSN 1521-4095. doi:10.1002/adma.202310104.

